# Spatial single cell multi-omics identifies divergent melanoma-cell states associated with metastatic outcomes in thin primary melanomas

**DOI:** 10.64898/2026.11.19.767941

**Authors:** Asier Antoranz, Madhavi Dipak Andhari, Jonathan Chui, Nikolina Dubroja, Pouya Nazari, Niels Vandermeulen, Katy Vandereyken, Jérémie Nsengimana, Canan Güvenç, Philip Georg Demaerel, Delphine Verstegen, Annemiek Leeman, Joanna Pozniak, Daniela Massi, Vincenzo De Giorgi, Anna Szumera-Ciećkiewicz, Piotr Rutkowski, Paweł Teterycz, Antien Mooyaart, Sara Valpione, Veerle Boecxstaens, Thierry Voet, Olivier Govaere, Francesca Bosisio

**Affiliations:** Translational Cell & Tissue Research, Department of Imaging & Pathology, KU Leuven, Leuven, Belgium; KU Leuven Institute for Single Cell Omics (LISCO), KU Leuven, Leuven, Belgium; Laboratory for Reproductive Genomics, Department of Human Genetics, KU Leuven, Leuven, Belgium; Population Health Sciences Institute, NIHR Biomedical Research Centre, Newcastle University, Newcastle upon Tyne, United Kingdom; Laboratory of Dermatology, Department of Oncology, KU Leuven, Leuven, Belgium; Department of Dermatology, University Hospitals Leuven, Leuven, Belgium; Laboratory for Molecular Cancer Biology, VIB–KU Leuven Center for Cancer Biology, KU Leuven, Leuven, Belgium; Department of Health Sciences, University of Florence, Florence, Italy; Histopathology and Molecular Diagnostics, Careggi University Hospital, Florence, Italy; Department of Molecular Pathobiology, New York University – College of Dentistry, New York, USA; Section of Dermatology, Department of Health Sciences, University of Florence, Florence, Italy; Department of Cancer Pathomorphology, Maria Skłodowska-Curie National Research Institute of Oncology, Warsaw, Poland; Biobank, Maria Skłodowska-Curie National Research Institute of Oncology, 02-781 Warsaw, Poland; Department of Soft Tissue/Bone Sarcoma and Melanoma, Maria Skłodowska-Curie National Research Institute of Oncology, 02-781 Warsaw, Poland; Department of Pathology and Clinical Bioinformatics, Erasmus MC, University Medical Center Rotterdam, Rotterdam, the Netherlands; Division of Cancer Sciences, University of Manchester, Manchester, United Kingdom; The Christie NHS Foundation Trust, Manchester, United Kingdom; Cancer Research UK National Biomarker Centre, The University of Manchester, Manchester, United Kingdom; Surgical Oncology, Department of Oncology, KU Leuven, Leuven, Belgium; Department of Surgical Oncology, University Hospitals Leuven, Leuven, Belgium; Department of Pathology, University Hospitals Leuven, Leuven, Belgium

## Abstract

Thin melanomas (Breslow thickness ≤1 mm) generally have favorable outcomes, yet some metastasise, highlighting the need for biomarkers of progression. Here, we combined same-section single-cell spatial transcriptomics and proteomics to identify seven melanoma-cell states organized along two inferred trajectories sharing a common trunk, extending from an in situ epithelial-adjacent state through developmental stress to melanocytic differentiation before diverging into two lineages. Lineage 1 progressed through an IFN-associated stress intermediate state towards mesenchymal-invasive cells enriched in metastasizing tumors and associated with immune-hot neighborhoods, extracellular-matrix remodeling and angiogenesis, potentially supporting vascular mimicry and metastatic dissemination. Lineage 2 terminated in cohesive nevus-like cells retaining melanocytic features within relatively immune-cold neighborhoods and enriched in non-metastasizing tumors. We confirmed these associations using H&E-inferred spatial transcriptomics in an international multicentric cohort and independent bulk transcriptomic datasets in primary melanomas, linking spatially organized melanoma-cell programs to metastatic outcome and identifying potential biomarkers of progression in thin melanoma.

**Statement of significance:** Spatial multi-omics identifies divergent melanoma-cell trajectories associated with metastatic outcome in thin melanomas. Their distinct microenvironments, protein profiles and transcriptional programs connect cellular heterogeneity to clinical outcome. These associations are supported by independent histological and bulk transcriptomic cohorts of primary melanomas, highlighting the potential of melanoma-cell states for risk stratification.

**Graphical Abstract:** 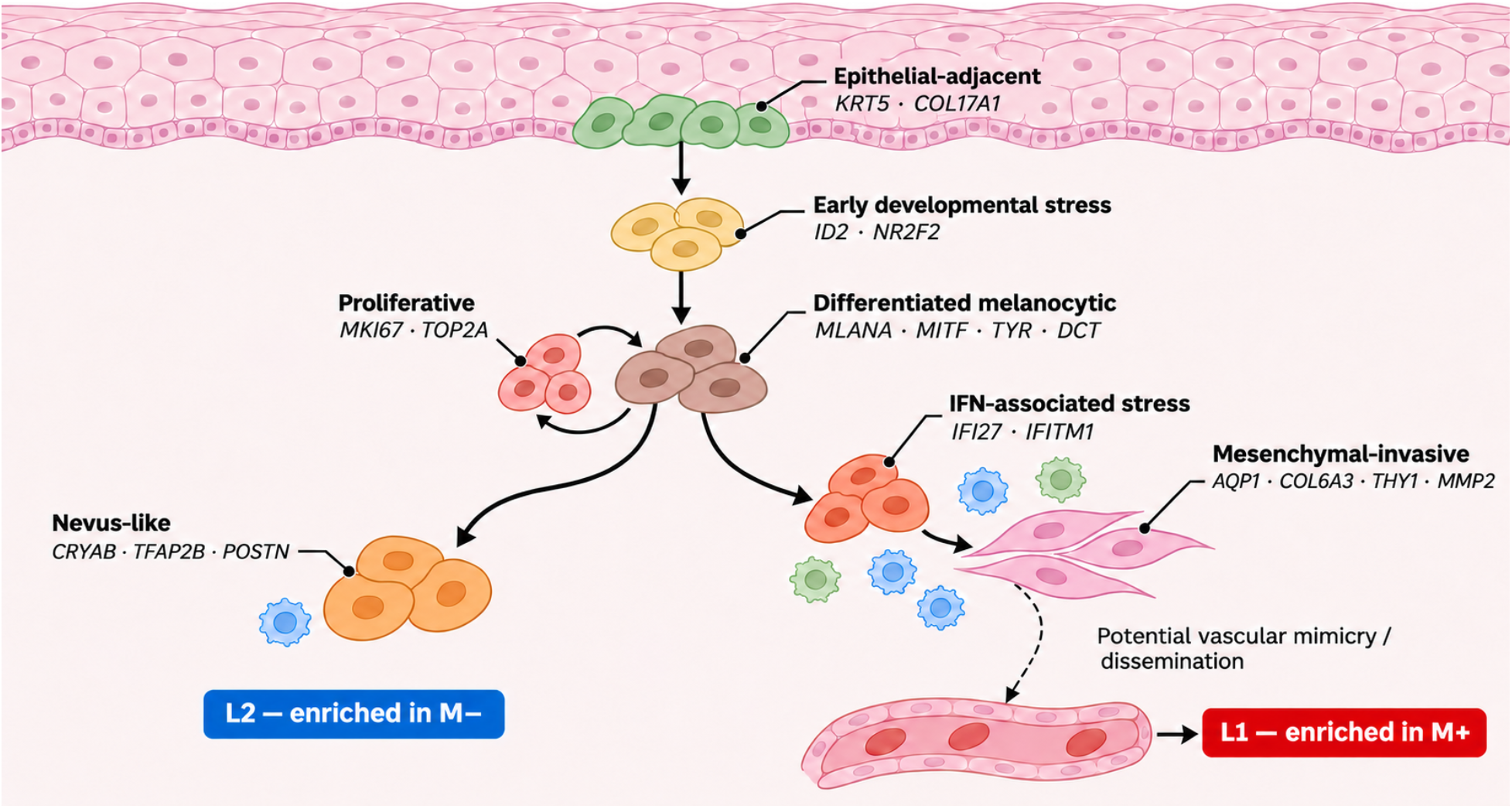

In this study, we combined spatial transcriptomics and multiplexed protein imaging to characterize melanoma cell heterogeneity and its spatial environment in thin primary cutaneous melanoma and linked it to metastatic outcome. We describe two trajectories that recapitulate the spatial transition from in-situ to invasive melanoma: epithelial-adjacent cells above the dermal–epidermal junction progressed through developmental stress towards a differentiated melanocytic state in the invasive component. From this common trunk, two trajectories extended deeper into the tumor: one towards mesenchymal-invasive cells associated with immune-hot neighborhoods and metastatic outcome; and another one towards nevus-like cells associated with relatively immune-cold neighborhoods and non-metastatic outcome. Proliferation represented an orthogonal program in line with historical dichotomic classification of melanoma cell states (invasive vs proliferative).

## Introduction

Breslow thickness is central to melanoma prognosis and clinical management, with increasing tumor thickness associated with greater metastatic risk (De Giorgi, et al., 2024). Melanomas with a Breslow thickness ≤1 mm (thin melanomas) generally have favorable outcomes and, when localized, are managed primarily by surgical excision (Amaral et al., 2025). However, some patients develop metastatic disease despite their limited primary tumor thickness. In one cohort of patients with metastatic melanoma, at least 10.23% were initially diagnosed with early-stage melanoma (Ertekin et al., 2021). In a population-based cohort of 144,447 patients diagnosed with early-stage melanoma, the cumulative incidence of melanoma-related death reached 6.3% at 20 years (Lo et al., 2025). Identifying patients at increased risk of progression could help guide intensified surveillance mechanisms or additional treatment strategies.

Why some thin melanomas develop metastases while others do not remains unclear. Investigating this question is challenging because the small size of these lesions and their routine processing as formalin-fixed, paraffin-embedded (FFPE) tissue for diagnosis limit access to fresh material for conventional single-cell RNA sequencing (Slyper et al., 2020; Wang et al., 2024b). Bulk profiling remains an alternative, but signals from small populations of invasive melanoma cells can be diluted by those from surrounding immune and stromal cells (Tirosh et al., 2016). Single-nucleus RNA sequencing enables profiling of archival tissue (Wang et al., 2024b), but small invasive populations may be sparsely sampled, and tissue dissociation removes the spatial context needed to distinguish invasive from in situ melanoma cells (Nirmal et al., 2022). Conventional Immunohistochemistry (IHC) preserves tissue architecture and works with archival material, but typically measures one marker at the time, limiting the range of cell types and states that can be characterized (Lin et al., 2018).

These technological limitations are reflected in the existing biomarker literature that have identified associations with outcome without fully resolving this cellular and spatial heterogeneity. Previous IHC studies have examined peritumoral CD8⁺ and FOXP3⁺ lymphocytes in thin melanomas (Sabbatino et al., 2020) and a seven-marker immunohistochemical signature in stage I–IIA disease (Meyer et al., 2023). Bulk transcriptomic analysis has also linked higher *TERT* expression in metastases to lower thickness of the originating primary melanoma (Kuhn et al., 2023). Gene-expression tests have also been developed, although their routine use to guide clinical management is not currently recommended (Bartlett et al., 2025). These studies identify biological associations but do not resolve the diversity and spatial organization of melanoma-cell states within thin melanomas.

Single-cell profiling of melanoma has identified coexisting differentiation programs, including melanocytic, intermediate and mesenchymal-like states, some of which have been linked to tumor growth and metastatic initiation (Tirosh et al., 2016; Wouters et al., 2020; Karras et al., 2022). However, these findings derive mainly from metastatic lesions, melanoma cell lines and experimental models. Spatial transcriptomics and proteomics now allow melanoma-cell heterogeneity to be studied directly in archival primary lesions while preserving tissue architecture. Using multiplexed imaging and regional transcriptomics, Nirmal et al. mapped cellular neighborhoods and immune organization across histological stages of primary melanoma (Nirmal et al., 2022). Love et al. used single-cell spatial transcriptomics to compare benign nevi, primary melanomas and metastases, identifying progression-associated programs and prognostic signatures (Love et al., 2024). Despite these advances, melanoma-cell heterogeneity in thin melanomas and its relationship to subsequent metastatic outcome remains poorly characterized.

Here, we investigated melanoma-cell heterogeneity, spatial organization and microenvironmental context in primary thin melanomas from patients who subsequently developed metastases (M+) and matched patients who remained metastasis-free (M-). We combined Xenium single-cell spatial transcriptomics with MILAN multiplexed protein imaging on the same tissue sections and Visium CytAssist transcriptomics on a serial section. We identified seven melanoma cell states organized along two inferred trajectories sharing a common trunk: one extending through an IFN-associated stress state towards a mesenchymal-invasive state, enriched in M+ melanomas, and another towards a nevus-like state, enriched in M-melanomas. We further assessed these associations using spatial gene expression inferred from H&E images in an independent international multicenter cohort and two publicly available bulk transcriptomic datasets. These validation cohorts included primary melanomas without restriction to Breslow thickness ≤1 mm. Together, these analyses link melanoma-cell states and their tissue organization to subsequent metastatic outcome in thin melanomas and support their clinical relevance in broader primary-melanoma cohorts.

## Results

### A multimodal spatial atlas of thin melanomas

To investigate cellular features associated with subsequent metastatic outcome, we assembled a discovery cohort of ten thin melanomas (5 M+, 5 M−; Supplementary Table 1). Xenium spatial transcriptomics measured 260 genes across 21 tissue sections, with repeated sections from each tumor (one tumor had 3 sections). Six tumors were additionally profiled using MILAN multiplexed protein imaging (Bolognesi et al., 2017) on a 42-marker panel on the same sections and whole-transcriptome Visium on serial sections (Fig. 1A–C). We registered the images from the three modalities using VALIS (Gatenbee et al., 2023), placing both RNA expression modalities and protein expression in a common spatial coordinate system. This linked RNA and protein measurements in 217,798 matched cell pairs and placed these measurements within the broader transcriptional context of the Visium spots (Fig. 1E-G; Supplementary Fig. 1A–I).

**Figure 1.**
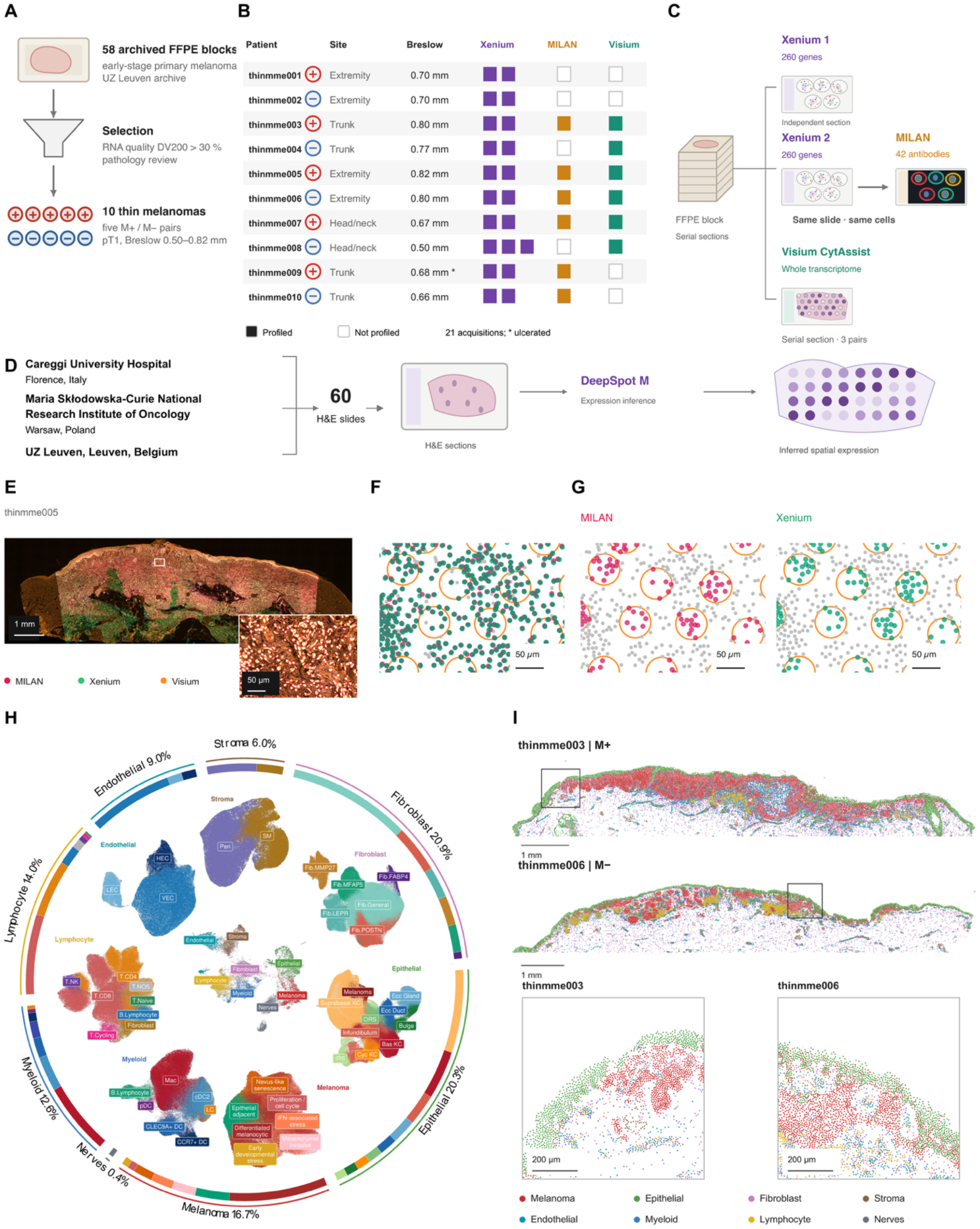
Study design and multimodal spatial atlas of thin melanomas. **A)** Selection of ten thin primary melanomas from 58 archived formalin-fixed, paraffin-embedded tissue blocks. Five patients who subsequently developed metastases (M+) were matched to five patients who did not during follow-up (M−) by the main clinicopathological variables (Breslow thickness, anatomical site, age, sex, ulceration). **B)** Clinical characteristics and assay availability for each tumor. **C)** Tissue-section workffow combining Xenium spatial transcriptomics using a 260-gene panel, subsequent MILAN multiplexed imaging using 42 antibodies on the same section, and whole-transcriptome Visium profiling on serial sections in subsets of the cohort. **D)** Independent multicenter validation cohort comprising 60 slides from 58 patients collected at three hospitals. Spatial gene expression was inferred from hematoxylin and eosin (H&E) images using DeepSpot-M. **E)** Representative registration of Xenium, MILAN and Visium measurements, with an enlarged view of the indicated region. **F)** Representative pairing of cells between registered Xenium and MILAN images. **G)** Assignment of MILAN and Xenium cells to Visium spots. **H)** UMAP overview of the Xenium atlas, showing cell types and their subtypes. Colored ring segments indicate subtype proportions. The outer color bands and labels indicate the eight broad lineages and their percentages. **I)** Representative spatial distributions of the eight broad lineages in M+ and M− melanomas, with enlarged views of the indicated regions.

After QC filtering and normalization, the Xenium atlas contained 1,365,249 cells from 19 acquisitions (2 acquisitions were dropped during QC) (Supplementary Fig. 2; Methods). We identified eight broad cellular lineages based on marker expression and spatial localization: epithelial (20.3%; *KRT5*, *KRT15*), endothelial (9.0%; *CDH5*, *VWF*), fibroblast (20.9%; *FBLN1*, *PDGFRA*), lymphocyte (14.0%; *CD3D*, *CD3E*), melanoma (16.7%; *MLANA*, *PMEL*), myeloid (12.6%; *LYZ*, *CD68*), nerve (0.4%; *PLP1*, *S100B*) and stromal (6.0%; *ACTA2*, *TAGLN*) populations (Supplementary Fig. 3; Fig. 1H–I).

To resolve non-melanoma subtypes, we analyzed each compartment separately, combining SingleR reference-based annotation (Aran et al., 2019) with marker-based annotation. For label transfer we used a publicly available integrated single-cell RNA-sequencing atlas of human skin (Restrepo et al., 2026). We first assessed whether the reference populations could be resolved using the 260 gene panel measured by Xenium (Supplementary Table 2). Populations that could not be reliably separated were merged, and marker-based annotation was used where reference mapping was insufficient. The final subtype annotations were supported by their corresponding marker-expression profiles and checked against their locations within the tissue (Supplementary Fig. 4A–F; Supplementary Table 3).

### Melanoma-cell metaprograms define divergent trajectories and spatial organization in thin melanomas

To characterize melanoma-cell heterogeneity, we identified groups of genes that were recurrently detected together within melanoma cells. Gene co-detection analysis retained a connected network of 122 genes and identified seven modules (Fig. 2A; Supplementary Fig. 5A–D; Supplementary Table 3). We scored these programs in individual melanoma cells and assigned each cell to its highest-scoring program (Fig. 2B). These metaprograms were annotated using their constituent genes, their overlap with published transcriptional signatures, and their spatial location (Fig. 2C; Supplementary Table 3). Three programs corresponded to established melanoma phenotypes. The “differentiated melanocytic” program contained pigmentation-associated genes, including *TYR*, *DCT* and *MITF*, and overlapped with published melanocytic and MITF-target signatures (Tirosh et al., 2016; Rambow et al., 2018; Wouters et al., 2020). The “proliferative” program contained cell-cycle genes, including *MKI67*, *TOP2A* and *CDK1*, and overlapped with published melanoma cell-cycle signatures. The “mesenchymal-invasive” program contained extracellular-matrix and mesenchymal-associated genes, including *COL6A3*, *MMP2* and *THY1*, and overlapped with published mesenchymal-like melanoma signatures and the Hallmark epithelial-to-mesenchymal transition gene set (Karras et al., 2022; Pozniak et al., 2024).

**Figure 2.**
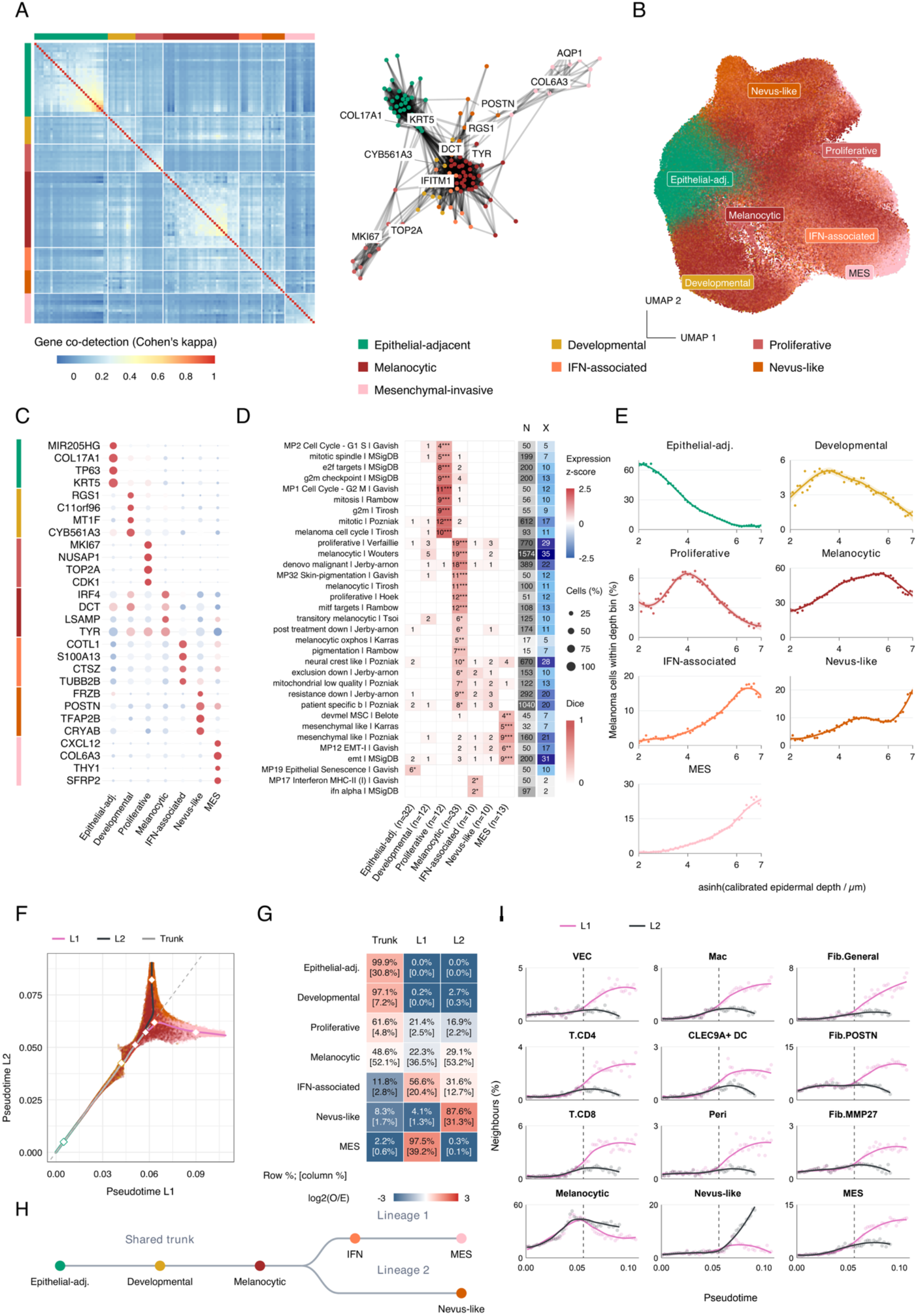
Melanoma metaprograms, inferred trajectories and spatial organization in thin melanomas. **A)** Gene co-detection matrix and network defining seven metaprograms. **B)** UMAP of melanoma cells colored by assigned metaprogram. **C)** Dotplot showing top four genes per program selected by log₂ fold change. **D)** Overlap with published signatures. Column labels report metaprogram gene counts; N, X, and n indicate total reference-signature genes, those represented in the Xenium panel, and metaprogram signature size, respectively. Asterisks indicate Benjamini–Hochberg-adjusted hypergeometric P values: * <0.05, ** <0.01 and *** <0.001. **E)** Metaprogram proportions among melanoma cells within epidermal-depth bins. **F)** Two-dimensional representation of lineage-specific pseudotimes, colored by metaprogram. Curves indicate the shared trunk and two inferred lineages (L1,L2); diamonds indicate median metaprogram positions. The dashed diagonal marks equal pseudotime values. **G)** Correspondence between metaprograms and trajectory assignments. Values indicate percentages within each metaprogram, with percentages within each trajectory class in brackets. **H)** Schematic summary of the inferred shared trunk and divergent lineages. **I)** Local cellular neighborhood composition along each lineage, using their 25 nearest non-self neighbors.

Two additional programs showed correspondence with broader cancer-cell programs. The “epithelial-adjacent” program contained epithelial and basal-cell-associated genes, including *KRT5*, *KRT15* and *COL17A1*, and overlapped with the pan-cancer epithelial-senescence metaprogram (Gavish et al., 2023). The “IFN-associated stress” program contained interferon-responsive genes, including *IFI27* and *IFITM1*, and overlapped with the Hallmark interferon-α response and a pan-cancer interferon/MHC-II metaprogram (Gavish et al., 2023) (Supplementary Fig. 5E–F).

The remaining two programs (5 and 6) showed no significant overlap with the tested public signatures, and their annotations were therefore based on the genes present in their signatures. Metaprogram 5 combined developmental regulators, including *ID2* and *NR2F2*, with the stress-associated gene *MT1F*, suggesting a program coupling developmental regulation with cellular stress (Martinsen and Bronner-Fraser, 1998; Davalos et al., 2023). We designated this program “early developmental-stress” (Fig. 2C; Supplementary Table 3). Module 6 was characterized by genes including *CRYAB*, *POSTN*, *TFAP2B* and *IGFBP7* (Fig. 2C; Supplementary Table 3). Although *TFAP2B* and *POSTN* were shared with published neural crest-like melanoma signatures, the overall overlap was not significant. The program also contained *IGFBP7*, whose expression has been reported in benign nevi (Wajapeyee et al., 2008, 2010). This motivated investigating whether this program resembled expression states found in nevi. We projected all seven metaprograms onto an independent bulk transcriptomic cohort including primary melanomas and benign melanocytic nevi (Kunz et al., 2018) (Supplementary Fig. 5G). Metaprogram 6 scores were higher in nevi than in melanomas (FDR < 0.001), supporting its designation as “nevus-like” (Supplementary Fig. 5H–I; Supplementary Table 3). Xenium transcripts show the spatial expression of metaprogram-associated genes alongside *MLANA*, *PMEL* and *S100B* in representative tissue regions (Supplementary Fig. 6).

We next checked the location of these melanoma metaprograms using the epidermal basal layer as spatial reference. Overall melanoma-cell abundance was highest near the epidermis and declined with increasing depth (Supplementary Fig. 7A). Across metaprograms epithelial-adjacent cells dominated near the epidermis and declined sharply with increasing depth. The developmental and proliferative programs peaked at relatively shallow depths, followed by the melanocytic program, whereas the IFN-associated, nevus-like and mesenchymal-invasive programs became more prominent at greater depths (Fig. 2E).

We next examined the relationships between the metaprograms using a diffusion-map embedding of their module scores. The proliferative program represented a different axis of heterogeneity (Supplementary Fig. 7B), and was excluded from model fitting, in line with previous studies (Liang et al., 2020; Ranek et al., 2024). We therefore inferred trajectories from the six remaining programs using Slingshot (Street et al., 2018), with the epithelial-adjacent state as the root being the most superficial melanoma metaprogram (Supplementary Fig. 7C). Trajectory inference identified a shared trunk extending from epithelial-adjacent through early developmental stress to differentiated melanocytic states, followed by two branches. Branch 1 (Lineage 1) extended through IFN-associated stress towards a mesenchymal-invasive state, whereas branch 2 (lineage 2) extended towards a nevus-like state (Fig. 2F,H; Supplementary Fig. 7D–J). Mean pseudotime increased with distance from the epidermal basal layer along both lineages (Spearman’s ρ = 0.99, P < 2.2 × 10⁻¹⁶ for both), linking the inferred trajectories to increasing tissue depth (Supplementary Fig. 7K). These trajectories extend the early distinction between proliferative and invasive melanoma states (Hoek et al., 2006, 2008), resolving two divergent routes within the invasive tumor compartment towards mesenchymal-invasive and nevus-like phenotypes.

To characterize their local environments, we quantified the cell types among the 25 nearest neighbors of each melanoma cell. Neighborhood composition differed across metaprograms: epithelial-adjacent cells were frequently surrounded by keratinocytes (45.9%) and other epithelial-adjacent melanoma cells (18.6%), whereas mesenchymal-invasive cells had higher contributions from immune (18.5%), fibroblast (16.4%) and vascular populations (3.6%) also when compared to other deep metaprograms such as nevus-like (8.6%, 9.4%, and 1.8% respectively). Melanocytic and nevus-like cells had neighborhoods dominated by melanoma cells (68.3% and 64.0% respectively) (Supplementary Fig. 8A).

Grouping cells by trajectory class similarly distinguished an epithelial-rich shared trunk, more immune- and stromal-rich neighborhoods around lineage 1 cells, and a larger melanoma-cell component around lineage 2 cells (Supplementary Fig. 8B). Analysis along pseudotime showed that these differences emerged predominantly after the fork: lineage 1 had higher fractions of vascular endothelial cells, macrophages, CD4⁺ and CD8⁺ T cells, dendritic cells, fibroblasts and pericytes, whereas lineage 2 retained higher fractions of melanocytic and nevus-like melanoma neighbors (Fig. 2I; Supplementary Fig. 8C).

### Cellular composition associated with metastatic outcome

Before comparing cellular composition between M+ and M− tumors, we assessed concordance across repeated Xenium acquisitions. Cellular composition was more similar between repeated sections from the same tumor than between sections from different tumors (permutation P < 0.0001; Supplementary Fig. 8D–G). Two acquisitions showed limited melanoma-cell recovery relative to other sections from the same tumor and were excluded from downstream analyses (Supplementary Fig. 8H). We pooled cell counts across the remaining 19 acquisitions to derive patient-level subtype proportions within each broad cellular compartment (Supplementary Table 4).

We then compared matched patient-level cell proportions between M+ and M− tumors using propeller (Phipson et al., 2022). Within the melanoma compartment, the nevus-like program accounted for a larger fraction of cells in M-compared to M+ (17.5% vs 3.3%; FDR = 0.012), with larger proportions in all five matched pairs. Differences also involved the lymphocyte and fibroblast compartments. CD8⁺ T cells constituted a larger fraction of lymphocytes in M-tumors (51.2% versus 44.1%; FDR = 0.0036). Within fibroblasts, M+ tumors had lower proportions of the *LEPR* subtype (8.7% versus 15.9%; FDR = 0.013) and higher proportions of the general fibroblast subtype (47.6% versus 39.2%; FDR = 0.041; Fig. 3A; Supplementary Fig. 9A; Supplementary Table 4). When melanoma cells were grouped by trajectory class, the mean contribution of lineage 2 was lower in M+ tumors (19.0% versus 45.0%), whereas lineage 1 contributed a larger fraction (30.3% versus 12.4%). However, neither comparison reached significance after multiple-testing correction (FDR = 0.056 and 0.115, respectively) (Fig. 3B; Supplementary Fig. 9A).

**Figure 3.**
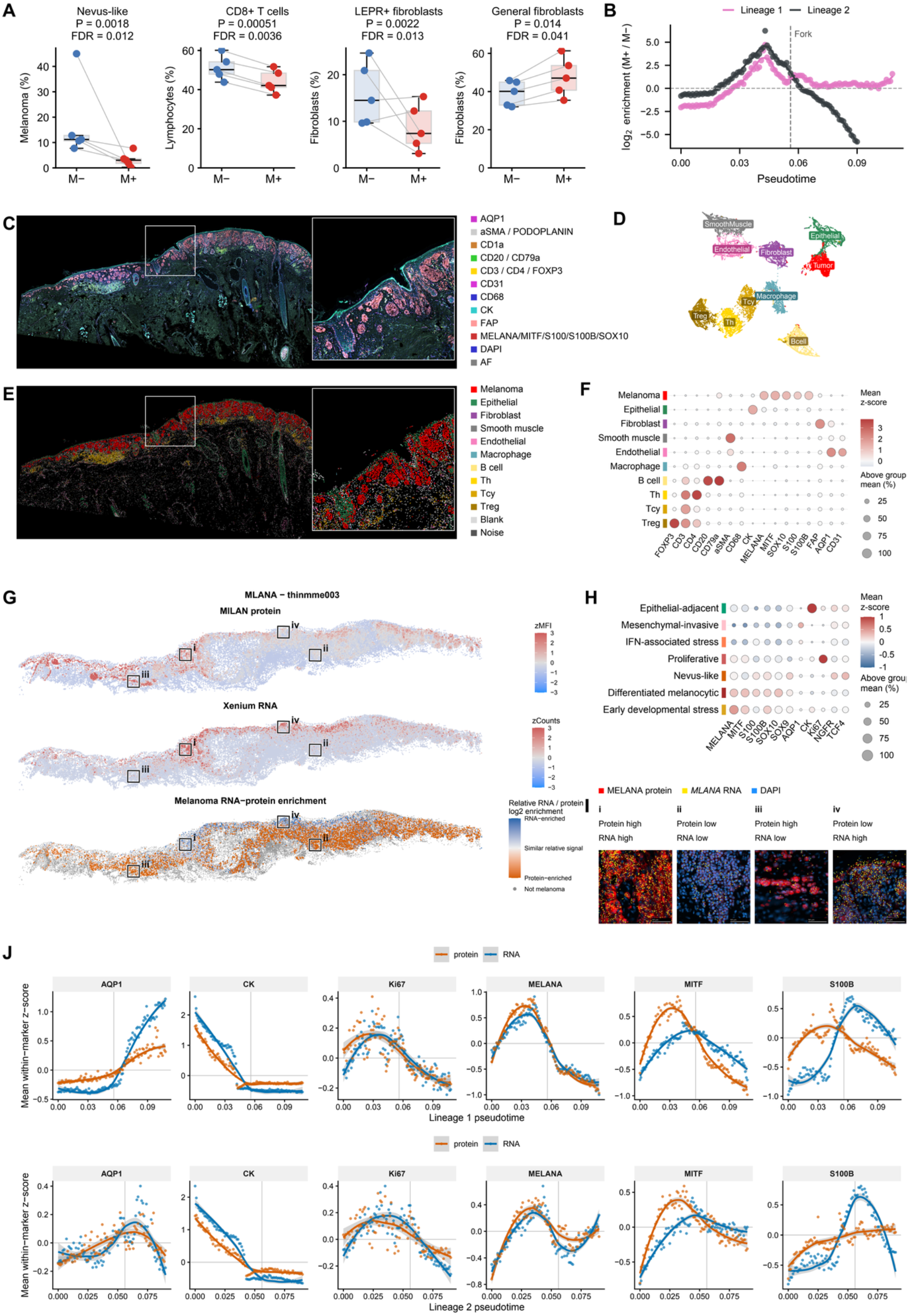
Cellular composition associated with metastatic outcome, protein profiling, and RNA–protein concordance. **A)** Patient-level proportions of selected cell populations in M+ vs M− patients, expressed within the indicated cellular compartment. P values and FDRs derive from paired propeller analyses, with Benjamini–Hochberg correction within each compartment. **B)** Relative enrichment of M+ versus M− melanoma cells along each trajectory, normalized to the overall M+/M− ratio within that trajectory. The shared trunk is included in both trajectories. **C)** Representative MILAN multiplex staining, with an enlarged view of the boxed region. **D)** UMAP of protein-profiled cells colored by annotated cell type. **E)** Digital reconstruction of the tissue in C, colored by cell type. **F)** Protein-marker profiles across annotated cell types. **G)** Spatial maps of MELANA protein and MLANA RNA in a representative example (thinmme003), displayed as modality-specific z-scores. The lower map shows relative RNA–protein enrichment in matched melanoma cells. **H)** Dotplot showing protein expression in melanoma metaprograms. **I)** Overlays of MELANA protein, MLANA transcript and DAPI images illustrating four combinations of relative protein and RNA abundance. **J)** RNA and protein profiles along Lineage 1 (upper row) and Lineage 2 (lower row), including the shared trunk. Signals were standardized separately for each marker and modality within each trajectory. Vertical lines mark the inferred fork.

### Same-section protein profiling of melanoma states

To examine the protein-level features of transcriptionally defined cell states, we performed MILAN multiplex immunofluorescence on six tissue sections previously profiled by Xenium, enabling paired RNA and protein measurements in individual spatially matched cells. Protein-based annotation resolved the major tumor, epithelial, immune and stromal populations (Fig. 3C–F). Among matched cells, general cell type agreement between modalities was high for melanoma, epithelial cells and lymphocytes (86.9%, 90.9% and 86.8%, respectively), but lower for fibroblasts and myeloid cells (18.8% and 46.1%). Agreement varied with the number of dedicated protein markers, although marker count alone did not explain the differences between lineages (Supplementary Fig. 9B–F; Supplementary Table 5).

Protein profiles supported selected metaprogram annotations, including higher cytokeratin expression in epithelial-adjacent cells and Ki67 expression in proliferative cells. Mesenchymal-invasive cells showed lower mean MELANA and MITF expression and higher relative AQP1 expression, whereas nevus-like cells showed higher relative expression of S100, S100B, SOX9, NGFR and TCF4 (Fig. 3H). Representative staining illustrated these marker combinations within their tissue context (Supplementary Fig. 10).

Matched RNA and protein measurements were positively correlated across the tested markers (Pearson r = 0.47-0.69), with variation between markers and samples (Supplementary Fig. 9G–J). Despite this overall correspondence, spatial overlays revealed local differences in relative RNA and protein abundance (Supplementary Fig. 11). These differences were often associated with tissue depth. Figure 3G-I illustrates this for thinmme003 where superficial melanoma regions contain higher relative *MLANA* RNA enrichment, whereas deeper regions contain greater relative MELANA protein enrichment. This spatial pattern suggests that transcriptional changes accompanying melanoma-cell dedifferentiation may precede changes in protein abundance, consistent with temporal delays between RNA and protein responses observed during cellular differentiation (Reimegård et al., 2021). Along the trajectories, in lineage 1, *MLANA* decreased and AQP1 increased at both RNA and protein levels (Fig. 3J), consistent with a transition towards an invasive phenotype, linking these molecular changes to the inferred trajectory.

### Whole transcriptome profiling of melanoma states with Visium

To extend the targeted Xenium and same-section protein analyses to broader transcriptional programs, we profiled serial sections from six tumors, comprising three matched M+/M− pairs, using Visium CytAssist. Quality filtering retained 7,652 spots across the six sections (Supplementary Fig. *12*A–C). RPCA Integration (Stuart et al., 2019) improved cross-section mixing, with kBET acceptance increasing from 0.8% to 20.0% (Büttner et al., 2019), (Supplementary Fig. *12*D–F). MAGIC imputation (van Dijk et al., 2018) increased spatial correspondence with registered Xenium pseudospots, with mean Lee’s L increasing from 0.147 to 0.202 (Lee, 2001) (Supplementary Fig. *12*G–I).

Xenium-guided deconvolution with cell2location (Kleshchevnikov et al., 2022) identified 1,590 melanoma-associated spots (Supplementary Fig. *13*A–B). Projection of the seven melanoma metaprograms and melanoma trajectories showed close agreement in patient-level composition with Xenium, with a median within-patient Spearman correlation of 0.978 (MAE = 0.020) for metaprograms and a pooled Spearman correlation of 0.915 (MAE = 0.055) across trajectories (Fig. 4A–B; Supplementary Fig. *13*C–F).

**Figure 4.**
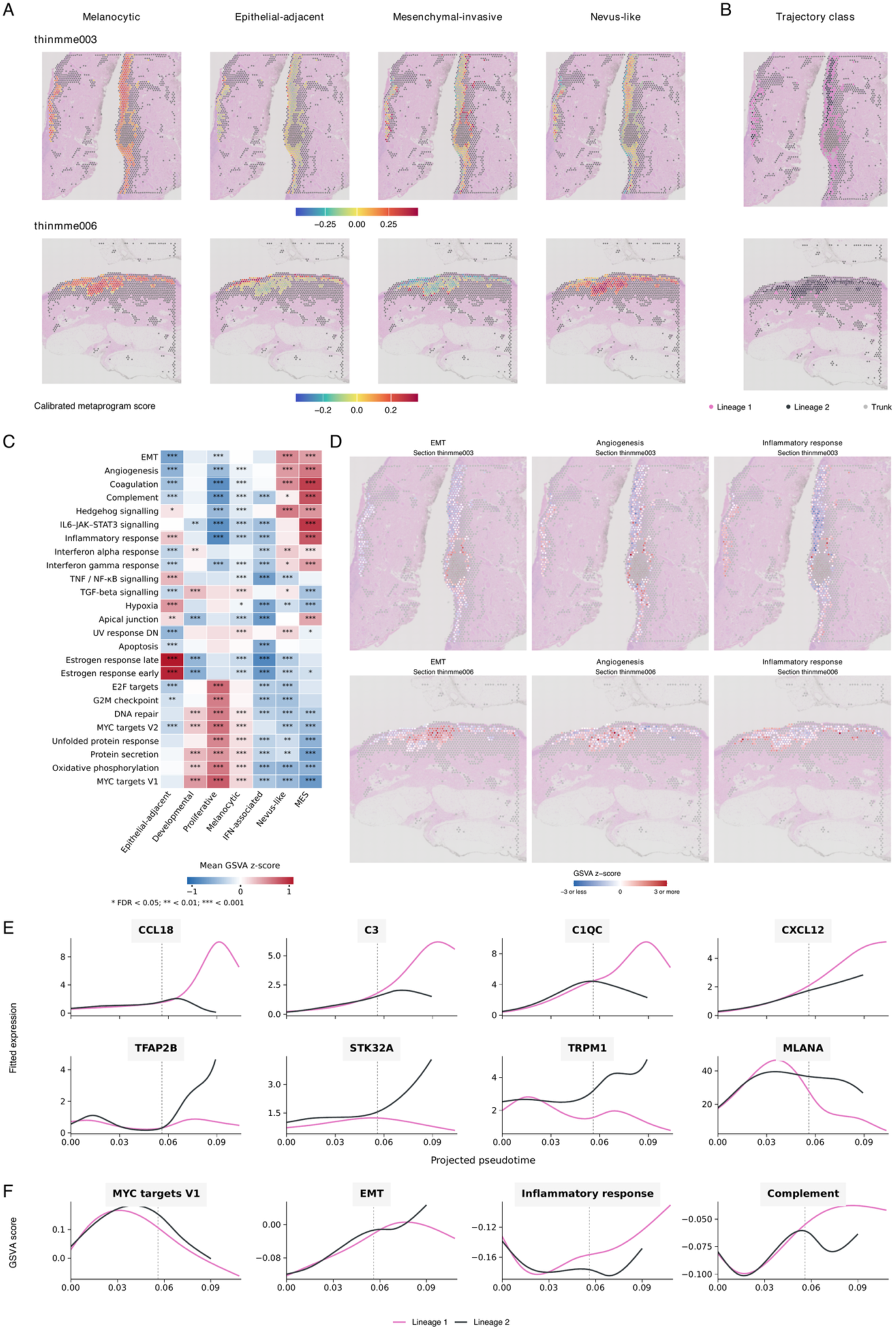
Visium profiling of melanoma metaprograms and trajectories. **A)** Spatial distributions of selected melanoma metaprogram scores in thinmme003 and thinmme006, overlaid on H&E images. **B)** Corresponding assignments to the shared trunk, Lineage 1 or Lineage 2. **C)** Mean GSVA z-score in spots assigned to each metaprogram in selected Hallmark. Asterisks indicate significance from Wilcoxon tests with Benjamini–Hochberg correction within each comparison (* FDR < 0.05; ** FDR < 0.01; *** FDR < 0.001). **D)** Spatial distributions of EMT, angiogenesis and inffammatory-response scores in the sections shown in A, displayed as z-scores. **E–F)** Fitted profiles of eight selected genes (E) and four Hallmark pathways (F) along projected pseudotime. The vertical dashed line indicates the fork.

Gene set variation analysis (GSVA) (Hänzelmann et al., 2013) of the 50 MSigDB Hallmark gene sets (Liberzon et al., 2015) revealed different pathway enrichment across metaprograms. Early developmental-stress, proliferative and differentiated-melanocytic regions showed higher scores for MYC targets, oxidative phosphorylation and protein secretion pathways, with E2F targets and G2M checkpoint particularly prominent in proliferative regions. Both mesenchymal-invasive and nevus-like regions showed elevated EMT and angiogenesis scores, whereas inflammatory-response and IL6–JAK–STAT3 signaling scores were more prominent in mesenchymal-invasive regions (all FDR < 0.0001) (Fig. 4C–D). Complete results for all 50 pathways are provided in Supplementary Table 6.

Whole-transcriptome and pathway analyses along inferred pseudotime further characterized the tissue environments associated with the two trajectories. Gene-expression profiles assessed using tradeSeq (Van den Berge et al., 2020) showed increasing *CCL18* expression along lineage 1 and higher *C3* and *C1QC* expression towards its endpoint. These changes were accompanied by higher inflammatory-response and complement pathway scores, particularly after the fork, consistent with the immune- and stroma-rich neighborhoods identified by Xenium, as *C3* expression has been reported in melanoma-associated fibroblasts and *C1QC* expression in macrophages (Davidson et al., 2020; Modak et al., 2022). In contrast, *TRPM1* expression increased towards late lineage 2, alongside greater retention of *MLANA* relative to lineage 1. As *TRPM1* is a direct MITF target associated with melanocytic differentiation, these findings supported the retained melanocytic features of the nevus-like endpoint (Miller et al., 2004). The estimated cellular composition helped contextualise these differences: mean relative melanoma weights were 26.2% in mesenchymal-invasive regions and 49.7% in nevus-like regions, with substantial fibroblast and myeloid contributions in mesenchymal-invasive regions. Thus, these profiles capture both melanoma-cell states and their associated tissue environments (Fig. 4E–F; Supplementary Fig. 13G; Supplementary Table 6).

### Validation of melanoma-state associations and assessment of clinical translatability

To assess whether the melanoma-state associations identified by spatial profiling could be recovered from routine histology, we analyzed H&E images from an international, multicenter primary melanoma cohort. This cohort was not limited to Breslow thickness < 1mm. The analysis included 58 patients with known metastatic outcome (35 M+ and 23 M−). DeepSpot-M (Nonchev et al., 2026) was used to predict patch-level expression of the Xenium-panel genes (257 in common with DeepSpot-M). After filtering melanoma-enriched spots, we scored all seven melanoma metaprograms and assigned each spot to its highest-scoring program (Fig. 5A–B; Supplementary Table 7). M-tumors showed a higher median nevus-like fraction than M+ tumors (18.2% versus 9.4%; FDR = 0.00042) and a lower median mesenchymal-invasive fraction (13.5% versus 29.4%; FDR = 0.00023) (Fig. 5C–D). The median MES/nevus-like ratio was 2.66 in M+ versus 0.63 in M− tumors, yielding a patient-level AUC of 0.843 (95% bootstrap CI, 0.737–0.930) (Fig. 5E).

**Figure 5.**
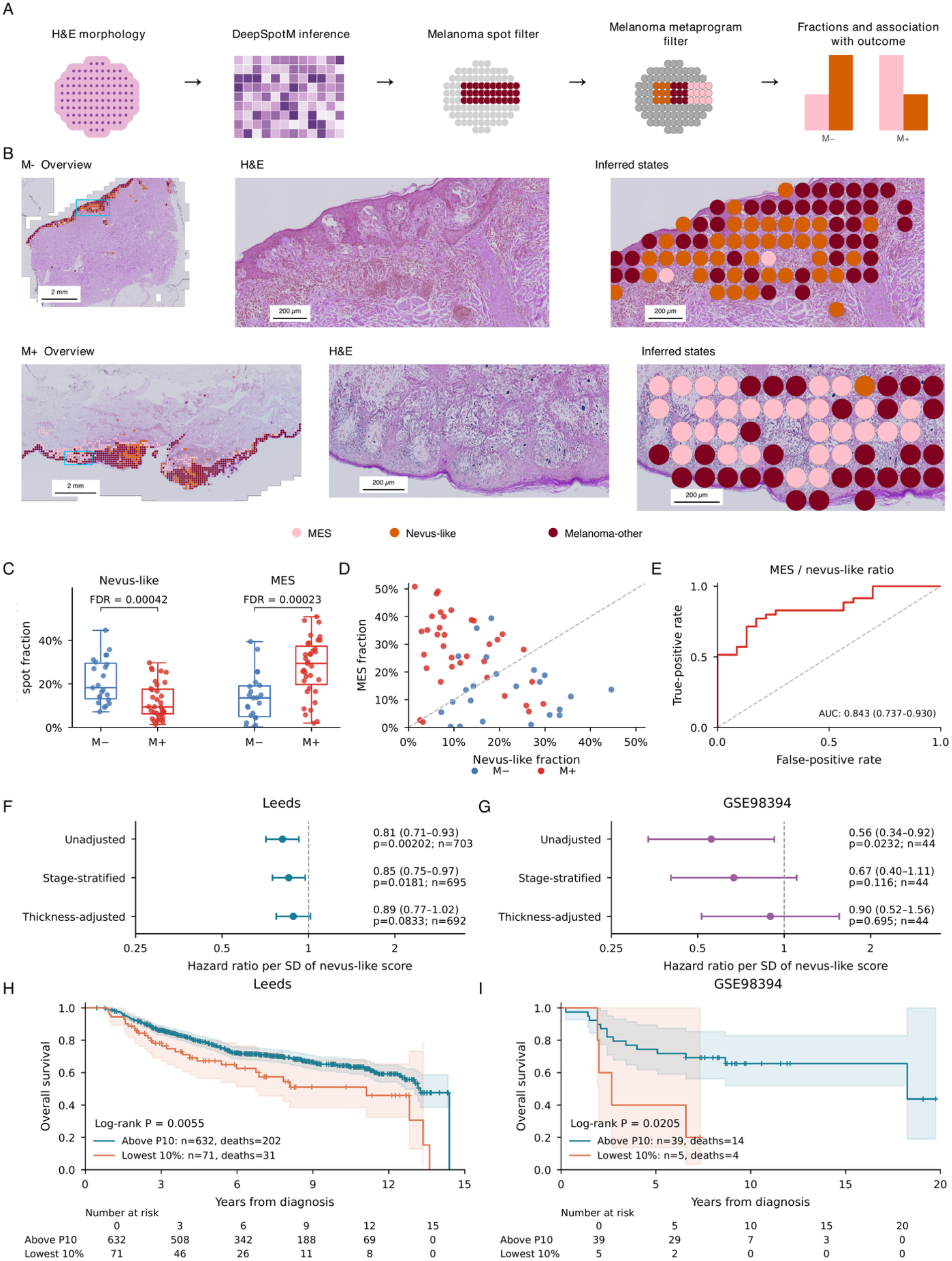
H&E-based inference of melanoma metaprograms and validation in bulk transcriptomic cohorts. **A)** Workffow for predicting gene expression from H&E using DeepSpot-M, identifying melanoma-enriched spots and assigning metaprograms. **B)** Representative M− and M+ sections, with assignments recalculated using the 257-gene Xenium panel. **C)** Nevus-like and MES fractions in 58 patients (35 M+, 23 M−). FDRs derive from two-sided Wilcoxon rank-sum tests with Benjamini–Hochberg correction. **D)** Joint patient-level fractions, colored by outcome (M+, red; M−, blue). **E)** ROC curve for the MES/nevus-like. Confidence intervals use 5,000 outcome-stratified bootstrap resamples. **F–G)** Associations between continuous nevus-like scores and overall survival in Leeds (F) and GSE98394 (G). **H–I)** Kaplan–Meier curves comparing the lowest 10% of nevus-like scores with the remaining samples.

We next examined whether melanoma metaprograms were associated with clinical outcome in additional primary-melanoma cohorts using bulk transcriptomics. Overall-survival analyses included 703 patients from the Leeds Melanoma Cohort (Nsengimana et al., 2018) and 44 from GSE98394 (Badal et al., 2017). We focused the bulk survival analyses on the nevus-like signature because MES-signature expression was more strongly attributable to non-melanoma populations. Reference-based estimates indicated median non-melanoma contributions of 68.8% in Leeds and 88.7% in GSE98394 for MES, compared with 20.9% and 51.2%, respectively, for nevus-like (Methods). Higher nevus-like scores were associated with lower mortality in both cohorts, with hazard ratios per standard-deviation increase of 0.81 in Leeds (95% CI, 0.71–0.93; P = 0.0020) and 0.56 in GSE98394 (95% CI, 0.34–0.92; P = 0.023). Although the statistical significance of the association was attenuated after accounting for stage (Leeds: HR = 0.85, P = 0.018; GSE98394: HR = 0.67, P = 0.116), or Breslow thickness (Leeds: HR = 0.89, P = 0.083; GSE98394: HR = 0.90, P = 0.695), higher nevus-like scores remained associated with lower estimated mortality across the adjusted models (Fig. 5F–G). To illustrate outcomes among patients with particularly low nevus-like scores, we compared the lowest-scoring decile with the remaining patients which showed poorer overall survival in both Leeds and GSE98394 (log-rank P = 0.0055 and P = 0.0205, respectively) (Fig. 5H–I) (Supplementary Table 7). These analyses support the clinical relevance of the melanoma-state differences identified by spatial profiling and their association with metastatic outcome and patient survival.

## Discussion

In this study, we combined spatial transcriptomics and multiplexed protein imaging to characterize melanoma-cell heterogeneity in thin primary melanomas with divergent metastatic outcomes. Our findings extend previous descriptions of melanoma-cell plasticity by identifying seven transcriptional metaprograms organized along two inferred trajectories, associated with distinct spatial microenvironments and clinical outcomes. Previous spatial studies mapped melanoma progression, immune organization and prognostic programs across histological stages and benign and malignant lesions (Nirmal et al., 2022; Love et al., 2024). Here, we focus on thin primary melanomas with different subsequent metastatic outcomes, linking melanoma-cell states to their spatial environments through same-section RNA and protein profiling.

Bulk transcriptomic studies initially described melanoma plasticity as switching between proliferative and invasive programs (Hoek et al., 2008; Verfaillie et al., 2015). Subsequent studies in scRNA-seq and bulk revealed multiple coexisting states and their underlying regulatory networks, developmental programs and immune-evasion mechanisms (Tirosh et al., 2016; Rambow et al., 2018; Tsoi et al., 2018; Jerby-Arnon et al., 2018; Wouters et al., 2020; Belote et al., 2021). Spatial analyses and lineage tracing further highlighted the influence of the tissue microenvironment on these states (Karras et al., 2022; Pozniak et al., 2024). How these states relate to metastatic outcome in thin primary melanomas remains less well characterized.

One of the most distinctive findings of our study is the identification of a nevus-like melanoma-cell metaprogram that has not been previously described. Although the melanocytic, proliferative and mesenchymal-invasive programs showed correspondence with established melanoma signatures, the nevus-like program remained distinct from published transcriptional references. Its biological identity was nevertheless supported by its enrichment in an independent dataset including primary melanoma and benign melanocytic nevi, its characteristic expression of *CRYAB* and *TFAP2B*, and its distinctive protein profile, including increased expression of S100, S100B, SOX9, NGFR and TCF4. Interestingly, the existence of melanoma cells displaying nevus-like morphological characteristics has long been recognized in dermatopathology through the phenomenon of melanoma pseudomaturation, whereby malignant melanocytes progressively decrease in size with increasing dermal depth, mimicking the maturation of benign melanocytic nevi (Ruhoy et al, 2000). This phenomenon highlights the remarkable morphological heterogeneity of melanoma cells and suggests that the acquisition of a nevus-like phenotype does not necessarily reflect benign differentiation.

An additional characteristic of the nevus-like population is its distinctive immune microenvironment. Both nevus-like and differentiated melanocytic melanoma cells were associated with neighborhoods predominantly composed of other melanoma cells, with relatively limited representation of immune populations. These two programs therefore appear to occupy the most immune-poor/cold tumor neighborhoods for the invasive component, although for potentially different biological reasons. In the differentiated melanocytic state, this observation may predominantly reflect the spatial organization of melanoma cells into nests, with limited intermixing of non-malignant cells. In contrast, the nevus-like program raises the possibility of an active immune-evasion phenotype despite its association with a more favorable clinical outcome. Of particular interest, nevus-like cells showed increased expression of TCF4 at the protein level. Previous work by Pozniak et al. (2024) identified TCF4 as a central regulator of melanoma dedifferentiation and resistance to immunotherapy, demonstrating its ability to suppress melanocytic differentiation, antigen-presentation programs and interferon signaling. These findings raise the possibility that TCF4 contributes to the immune-poor neighborhoods surrounding nevus-like cells. Whether this reflects a persistent, immune-evasive state, and how it relates to favorable outcomes in primary melanoma versus treatment resistance in metastatic disease, requires further investigation.

Additionally, Visium analyses revealed EMT and angiogenesis pathway enrichment in both nevus-like and mesenchymal-invasive regions, despite their distinct melanocytic and inflammatory profiles. However, the expression patterns of angiogenesis-associated genes differed between these regions: genes involved in vessel-formation signaling (*NRP1*, *FGFR1*, *VEGFA*) were more highly expressed in mesenchymal-invasive regions, whereas genes involved in extracellular-matrix organization and remodeling (*POSTN*, *COL3A1*, *COL5A2*) were more highly expressed in nevus-like regions. Together with the protein and spatial findings, these patterns raise the hypothesis that mesenchymal-invasive cells acquire vascular-associated properties, potentially contributing to vasculogenic mimicry and dissemination (Maniotis et al., 1999; Dunleavey et al., 2014), while nevus-like cells may additionally evade immune elimination through extracellular-matrix remodeling. Nevertheless, inferred trajectories do not establish actual cell-state transitions, and the proposed relationship between TCF4, the nevus-like phenotype and immune evasion remains to be functionally demonstrated.

Inferred molecular trajectories require cautious interpretation in cancer, where transcriptional similarities do not necessarily reflect actual cell-state transitions (Tirosh, 2026). This limitation is particularly relevant to dissociation-based scRNA-seq, where spatial information is lost. Spatial profiling provides additional evidence by locating cells along the inferred trajectory within the tissue. Here, the correspondence between pseudotime and tumor depth provides spatial support for the inferred ordering. We therefore interpret these trajectories as relationships among coexisting melanoma-cell states, rather than fixed, unidirectional paths of progression.

Because spatial biology and histopathology provide a snapshot of tumor progression at the time of surgical excision, enrichment of nevus-like melanoma cells may indicate a tumor state predominantly characterized by immune evasion rather than invasion, whereas a higher proportion of mesenchymal-invasive cells may reflect a tumor actively undergoing metastatic dissemination. We therefore propose the mesenchymal-invasive/nevus-like ratio at the time of excision as a potential predictor of the patient’s subsequent metastatic outcome. The main limitation of this study is the small discovery cohort, highlighting the need for validation in larger prospective studies. We demonstrated that the outcome-associated ratio of melanoma-cell states can also be recovered from H&E images, suggesting its potential applicability to routine diagnostic material. Although external H&E-based analyses supported the generalisability of the findings, computationally inferred gene expression cannot replace direct molecular measurements. These predictions may reflect melanoma-cell morphology, immune and stromal composition, and tissue architecture. Because melanoma states and their neighborhoods are associated in our spatial data, their contributions to the H&E predictions cannot be fully separated. Additionally, higher nevus-like scores were also associated with improved overall survival in two bulk transcriptomic cohorts. These findings support the exploration of different routes towards clinical implementation, although differences in cellular composition, disease stage, tumor thickness and other clinical characteristics complicate the interpretation of bulk transcriptomic results. Future studies should compare H&E-based inference, bulk transcriptomic scoring and reduced multiplexed panels for clinical risk stratification.

Spatially matched Xenium and MILAN analyses revealed important discrepancies between RNA and protein expression at the single-cell level. These differences may arise from transcriptional and post-transcriptional regulation, temporal delays between transcription and protein synthesis, protein turnover or technical variability, highlighting the complementary rather than interchangeable nature of the two modalities. Differences in subcellular RNA and protein localization, together with partial sampling of cells in tissue sections, may also contribute to the observed discordance (Adekunle and Wang, 2020). These differences caution against translating transcriptional metaprograms directly into protein markers. Clinical implementation will require reduced, validated RNA–protein panels, reproducibility across centers and prospective assessment of their added value beyond established clinicopathological factors, particularly in thin melanomas. An important next question is whether patients with an elevated predicted risk should undergo closer surveillance or could benefit from adjuvant immunotherapy before metastatic disease develops. Addressing this question will require clinical trials evaluating both treatment benefits and toxicity in patients selected on the basis of these biomarkers.

## Materials and methods

### Cohort description

Two cohorts were assembled: a discovery cohort from UZ Leuven for spatial transcriptomic and protein profiling, and an international, multicenter EORTC validation cohort for which only diagnostic hematoxylin and eosin (H&E) slides and clinical metadata were available.

For the discovery cohort, formalin-fixed, paraffin-embedded (FFPE) tissue blocks from 58 patients with thin melanoma were retrieved from the archives of UZ Leuven. RNA quality was assessed using the DV200 metric (Matsubara et al., 2020), with a DV200 >30% specified as the inclusion criterion for spatial transcriptomic profiling. All ten selected discovery samples exceeded this threshold (DV200 range, 31– 85%; Supplementary Table 1). From this pool, a dermatopathologist (FMB) selected ten representative thin melanoma cases: five from patients who later developed metastasis (M+) and five from patients with no record of metastasis development (M−). The M+ and M− cases were matched for key prognostic and clinicopathological features: Breslow thickness, patient age, ulceration status, tumor location, and histopathologic subtype. One pair did not match for ulceration. For each of these ten selected cases, tissue sections were cut from the same FFPE block for Xenium spatial transcriptomic profiling (10x Genomics). In six cases, MILAN (Bolognesi et al., 2017) was subsequently performed on the same section used for Xenium. Serial sections from six cases, comprising three matched M+/M− pairs, were processed using Visium CytAssist (10x Genomics).

The validation cohort included 60 diagnostic H&E slides from 58 patients (35 M+ and 23 M−), collected at Careggi University Hospital, Florence (7 slides), the Maria Skłodowska-Curie National Research Institute of Oncology, Warsaw (22 slides), and UZ Leuven (31 slides). The slides comprised 35 M+ and 25 M− specimens; two M− patients contributed two slides each. Breslow thickness ranged from 0.20 to 2.60 mm. Relevant clinicopathological characteristics for both cohorts are provided in Supplementary Table 1.

The study was approved by the Ethics Committee of UZ/KU Leuven (S61610). The Leeds Melanoma Cohort approvals were MREC1/03/57 and PIAG3-09(d)/2003.

### Data acquisition

#### Targeted Single-cell Spatial Transcriptomics with Xenium

Human skin ready-to-use panel (260 genes) was used to profile the explorative multiomics cohort. Two sections from each tumor were analyzed, with an additional section analyzed for thinmme008, yielding 21 acquisitions. Samples were analyzed according to the following demonstrated protocols, supplied by 10x Genomics. Any deviations from these demonstrated protocols are mentioned below.

1. CG000578 Rev. D (*B1*): Xenium In Situ for FFPE – Tissue Preparation Guide
2. CG000580 Rev. E (*B1*): Xenium In Situ for FFPE – Deparaffinization & Decrosslinking
3. CG000578 Rev. F (*B2*): Xenium In Situ - FFPE Tissue Preparation Handbook
4. CG000581 Rev. C (*B1*) and Rev. F (*B2)*: Xenium in Situ for Fresh Frozen Tissues – Fixation & Permeabilization
5. CG000582 Rev. H (*B1 + B2)*: Xenium in Situ Gene Expression User Guide
6. CG000584 Rev. F (*B1*) and Rev. K (*B2*): Xenium Analyzer

##### Sectioning

Xenium slides (10x Genomics, PN-3000941) were equilibrated to room temperature for 30 min prior to sectioning. Sectioning was performed on a microtome (HistoCore Multicut, Leica biosystems) at a thickness of 10 µm. After sectioning slides were dried overnight at room temperature, followed by a 3h bake at 42°C, and finally stored in a desiccator for a maximum of two weeks before proceeding with the remainder of the workflow.

The following paragraph is only applicable to samples processed in *batch 2.* Before sectioning, the paraffin on the FFPE block face was melted by placing the block face on a heat plate at 52°C for 5 mins. This was followed by an extended hydration of the block face for 30 min. Lastly the temperature of the water bath for floating the tissue sections was adjusted to 37°C.

##### Deparaffinization & Decrosslinking

Deparaffinization was initiated by incubating the slides at 60°C for 2 hours. The slides were then equilibrated to room temperature and immersed in the following solutions: twice 10 min in xylene, twice 3 min in 100% ethanol, twice 3 min in 96% ethanol, once 3 min in 70% ethanol and once in nuclease-free water. After deparaffinization, slides were inserted into Xenium cassettes, and 1x PBS was added to the cassette. 1x PBS was removed from the slide cassettes, followed by the addition of decrosslinking buffer and subsequent incubations of 30 min at 80°C and 10 min at 22°C in a thermocycler. Slides were retrieved from the thermocycler and washed thrice with PBS-T.

##### Probe Hybridization, Ligation & Amplification

PBS-T was removed from the cassette before the addition of Probe hybridization mix and a subsequent incubation for 18h in a thermocycler at 50 °C. Tissue was washed twice with PBS-T before the addition of post hybridization wash buffer (10x Genomics, PN-2000395) and incubation for 30 min in a thermocycler at 37°C. Tissue was washed thrice with PBS-T and Ligation mix was added before incubation for 2h in a thermocycler at 37°C. Amplification master mix was added after three PBS-T washes, and the slides were incubated for another 2h in a thermocycler at 30°C.

##### Autofluorescence Quenching & Nuclei Staining

Slides were washed thrice with TE buffer and thrice with 1x PBS. Tissue sections were incubated with diluted reducing agent B for 10 min at room temperature, followed by one wash with 70% ethanol and two washes with 100% ethanol. Slides were then incubated with autofluorescence quenching solution for 10 min at room temperature, followed by three 100% ethanol washes and drying on a thermocycler at 37°C for 5 min. Tissue was then rehydrated by the addition of 1x PBS and PBS-T subsequently. Nuclei were stained by the addition of nuclei staining buffer (10x Genomics, PN-2000762) for 1 min. Lastly the tissue sections were washed four times with PBS-T.

##### Xenium Analyzer

Samples were analyzed on a Xenium Analyzer instrument (10x Genomics, PN-1000481) running instrument software version 3.1.0.0 and Analysis version xenium-3.1.0.4 (batch 1), or instrument software version 4.0.1.4 and Analysis version xenium-4.0.1.0 (batch 2). Buffer and reagent compositions are provided in Supplementary Table 8.

#### Spatial Transcriptomics with Visium CytAssist

RNA quality of the FFPE blocks was determined prior to the Visium workflow. Sections of each block were collected in an Eppendorf tube, RNA was extracted using the RNeasy FFPE kit (Qiagen), and a DV200 score was ascertained using a BioAnalyzer system (Agilent).

Sample slide preparation was done according to Visium CytAssist Spatial Gene Expression for FFPE – Tissue Preparation Guide (CG000518 | Rev C, 10x Genomics). FFPE blocks were sectioned on a microtome (HistoCore Multicut, Leica biosystems) at a thickness of 10 µm and placed on SuperFrost Plus slides (631-0108, VWR). Tissue sections were dried overnight at room temperature before proceeding to further drying for 3 hours at 42°C.

Further processing of the samples was done according to Visium CytAssist Spatial Gene Expression for FFPE – Deparaffinization, HCE Staining, Imaging & Decrosslinking (CG000520 | Rev B, 10x Genomics) without any deviations. Samples retrieved from MILAN processing skipped the initial deparaffinization and started CG000520 Rev B at section 1.3, HCE staining. High resolution HCE images were generated using a custom system including a Nikon Eclipse Ni-E microscope body, ImagingSource DFK 33UX264 camera, and a Nikon plan apo λ 20x (0.75 NA) objective. qPCR results were obtained using a QuantStudio Pro 6 system (Applied Biosystems).

Final steps in sample processing and library generation were performed using the Visium FFPE Reagent Kit v2 according to Visium CytAssist Spatial Gene Expression Reagent Kits (CG000495 | Rev E, 10x Genomics). Libraries were sequenced on an Illumina NextSeq 2000 platform (paired-end, dual-indexing, 28 cycles Read 1, 10 cycles i7, 10 cycles i5, 50 cycles read 2).

High resolution images were manually aligned with the transcriptomic data using Loupe Browser version 6.5.0. Sequencing results were analyzed using Space Ranger version 3.1.1, and aligned to the GRCh38-2020-A reference genome.

#### Single-cell spatial proteomics with MILAN

Following Xenium processing, tissue sections from six cases were washed three times with PBS and subjected to antigen retrieval according to the MILAN protocol. Tissues were stained with a panel of 42 antibodies using iterative rounds of immunofluorescence staining, imaging and antibody stripping (Bolognesi et al., 2017). The antibody panel was designed to characterize the immune microenvironment and melanoma-cell phenotypic states (Supplementary Table 9). Antibodies were validated individually by comparing single-plex staining patterns with those reported in the Human Protein Atlas (Uhlén et al., 2015). Stripping efficiency was assessed by rescanning the tissue after antibody removal. Images were acquired at 10× magnification, with a resolution of 0.65 µm per pixel, using an Axio Scan.Z1 slide scanner (Zeiss, Germany) and saved in .czi format.

### Data Analysis

#### Cross-modal alignment (registration)

Images from tissue blocks profiled with more than one modality were spatially aligned before cross-modal analysis. Registration used DAPI images from Xenium and MILAN and H&E images from Visium. Images were resampled to a common pixel size of 0.6525 µm/pixel, oriented consistently, and preprocessed for intensity and contrast normalization. H&E images were converted to greyscale and inverted to facilitate alignment with fluorescence images. Registration was performed using VALIS (Gatenbee et al., 2023), with rigid alignment followed by non-rigid registration and Xenium as the reference image. Feature-detection settings were configured separately for each sample. The resulting transformations were applied to cell centroids and Visium spot centers to place them in a common coordinate system.

Cell segmentation and annotation were performed independently for each modality before matching (see the corresponding methods). Xenium and MILAN cells from the same tissue section were matched using their registered centroid coordinates. Matches were accepted when cells were reciprocal nearest neighbors, their separation was ≤5 µm, and the ratio of distances from the MILAN cell to its nearest and second-nearest Xenium cells was ≤0.7. These criteria provided one-to-one matches while excluding ambiguous assignments.

MILAN and Xenium cells were independently assigned to the nearest Visium spot when their registered centroids fell within its 27.5 µm radius. Cells outside this radius remained unassigned. Assignment rates were calculated among cells within 100/√3 ≈ 57.7 µm of the nearest spot center, corresponding to the maximum distance to a spot center within an ideal hexagonal grid with 100 µm center-to-center spacing. Registration, cell matching and spot assignment are illustrated in Supplementary Fig. 1.

#### Xenium analysis

##### Quality control and normalization

Xenium spatial transcriptomics data were processed in Seurat v5.4.0 (Hao et al., 2024). All samples and runs were merged into a single Seurat object. Quality control was based on the distribution of total transcript counts per cell (nCount_Xenium). A minimum threshold of 10 transcripts per cell was selected by inspecting the asinh-transformed count distribution, and cells below this threshold were excluded (Supplementary Fig. 2A–C).

Counts were normalized using Seurat’s LogNormalize method, with the scale factor set to the median total transcript count per cell after filtering (scale.factor = 101). This procedure divided each gene count by the cell’s total transcript count, multiplied it by the scale factor, and applied a log1p transformation. Normalized expression values were subsequently centered and scaled using ScaleData, with log1p-transformed total transcript counts and segmented cell area included as regression covariates to account for variation associated with count depth and cell size.

##### Dimensionality reduction and clustering of main cellular compartments

To define first-level cellular compartments in the Xenium dataset, we generated a balanced reference subset by randomly sampling up to 5,000 cells from each sample. Principal component analysis (PCA) was performed on the normalized data using all measured panel genes. UMAP dimensionality reduction (McInnes et al., 2018), nearest-neighbor graph construction, and graph-based clustering were then performed using the first 20 principal components. For clustering, resolution was set to 1. This yielded 29 initial clusters (Supplementary Figure 2E).

Potential acquisition- or sample-driven effects were assessed by visualizing the reference subset according to sample identity in the UMAP embedding (Supplementary Figure 2F). To quantify this, we computed the average silhouette width using sample identity as the grouping variable in PCA space (top 20 components) versus the same metric using cluster identity. Mean silhouette width was −0.065 for sample identity versus 0.154 for cluster identity (Wilcoxon rank-sum p < 2.2×10⁻¹⁶) (Supplementary Figure 2G), confirming the absence of dominant sample/run-driven clustering. Therefore, no explicit batch correction was applied before downstream clustering and annotation.

Clusters were transferred to the complete Xenium dataset using Seurat’s anchor-based label-transfer workflow (FindTransferAnchors/TransferData), computed on the complete gene panel and PCA-projected embeddings (Stuart et al., 2019). After projection, clusters were evaluated based on canonical marker expression (Supplementary Table 3), and expert pathological review (F.M.B.), visualizing cluster identities *in-situ* (Supplementary Figure 2H). This revealed cluster 16 as a mixture of undifferentiated melanoma cells and peripheral nerve (both neural-crest-derived). To resolve these, cluster-16 cells were extracted from the complete dataset and re-analyzed independently. To focus the embedding on informative markers within the cluster, feature selection was restricted to genes detected in at least 5% of cluster-16 cells. PCA, UMAP, nearest-neighbor graph construction and graph-based clustering (resolution 0.6) were then recomputed on this reduced feature set (Supplementary Figure 3A). Differentially expressed markers were calculated using Seurat’s FindAllMarkers function from where 10 neural and 10 melanoma markers were manually curated. The melanoma panel included: *PMEL*, *TYR*, *S100A13*, *MLANA*, *MITF*, *TYRP1*, *PCDH7*, *QPCT*, *EDNRB*, and *MYC*; while the neural panel included: *CRYAB*, *IGFBP5*, *NR2F2*, *TMEM176A*, *ITGA6*, *PLP1*, *DST*, *PTGDS*, *CXCL12*, *MYL9*. Each cell was scored (Seurat AddModuleScore) based on these two signatures (Supplementary Figure 3B–D) which cleanly identified subcluster 3 as peripheral nerve. This was further validated by mapping the classified cells *in situ* (Supplementary Figure 3E). After splitting nerves, the first annotation tier (Level 1) comprised of Melanoma, Epithelial, Fibroblast, Stroma, Endothelial, Myeloid, Lymphocyte, and Nerve populations.

##### Cell subtype annotation and reference mapping

Following broad cell-type classification, each major non-melanoma compartment (endothelial, myeloid, fibroblast, epithelial, stromal, and lymphocytic) was subtyped independently using a common workflow. The procedure uses reference-based label transfer as the default and falls back to de-novo marker annotation only where label transfer is demonstrably unreliable on the target panel, with every label ultimately confirmed against the observed Xenium expression and spatial location of the cells. For each lineage, cells were subset from the complete dataset and re-clustered using genes detected in at least 5% of the subset’s cells. PCA, nearest-neighbor graph construction (number of principal components selected for each lineage from the elbow plot: endothelial 1:5, myeloid 1:8, fibroblast 1:12, epithelial 1:10, lymphocyte 1:10, stroma 1:5) and graph-based clustering (resolution 0.8) were performed on the scaled data. Labels were transferred from a publicly available integrated skin scRNA-seq reference atlas (273,178 cells from 14 studies, detailed cell-type annotation with 41 classes) (Restrepo, P., et al., 2026) using SingleR (Aran D., et al., 2019), restricted to the subset of genes shared between the reference and the Xenium panel (255 of 260 panel genes) and the subset of cell types under that lineage.

Before mapping, we evaluated whether the atlas subtype labels are recoverable from the 255 common panel genes. Using the panel-restricted reference, we used 80% of the atlas to recover the held-out 20% (de.method = ‘wilcox’, aggr.ref = TRUE, fine.tune = TRUE). Performance was quantified by overall recovery (fraction of held-out cells correctly reassigned) and per-class recall, precision, and F1 (Supplementary Table 2). A transfer was accepted only when overall accuracy exceeded 0.8, and the transferred identities were consistent with the measured markers. When specific reference populations could not be separated on the measured markers, their labels were merged and the recovery reassessed. Marker-based cluster annotation was used when recovery remained below the threshold, or the reference categories did not adequately represent the observed populations.

**Endothelial (129,695 cells):** the atlas included high-endothelial cells (HEC), vascular endothelial cells (VEC), and lymphatic endothelial cells (LEC). Initial transfer over-estimated the number of HEC which were then merged with VEC achieving an overall recovery of 0.99. Clusters 6 and 8 were the only two with high levels of *SELE* and their transfer was manually overruled to HEC (Supplementary Figure 4A).

**Stroma (88,384 cells):** two stromal subtypes were present in the atlas: pericytes (Peri) and smooth muscle cells (SM). Overall recovery was below threshold (0.77) and SM had low F1 score (0.2), so the annotation was done de novo. The two states were then assigned per-cluster with *RGS5*+ and *NOTCH3*+ clusters being annotated as pericytes and *MYH11*+, *ACTA2*+, *TAGLN*+, *SYNPO2*+ and *MYL9*+ clusters annotated as SM (Vanlandewijck, M., et al., 2018) (Supplementary Figure 4B).

**Fibroblast (295,420 cells):** eight fibroblast subtypes were included in the atlas: papillary fibroblasts, reticular fibroblasts types I-III, perivascular fibroblasts types I-II, dermal papilla fibroblasts (DP) and dermal sheath fibroblasts (DS). Transfer of the atlas recovered at 0.74, and even though coarsening (merging the reticular and perivascular subtypes) lifted recovery to 0.81, the transfer was rejected: the atlas splits fibroblasts by dermal anatomy whereas the clusters are more functional and cancer-associated. Six states were therefore defined *de novo*: Fib general (*PI16*+, Buechler, M.B., et al., 2021), Fib *LEPR*+ (reticular/deep dermis, Buechler, M.B., et al., 2021), Fib *MFAP5*+ (*MFAP5*+ cancer associated fibroblasts, CAF, Wang, H., et al., 2024a), Fib *POSTN*+ (activated, ECM-remodeling, myofibroblastic cancer-associated fibroblast (myCAF), Liu, D., et al., 2026), Fib *FABP4*+ (lipid-associated, Fang C.L., et al., 2016), and Fib *MMP27*+ without a clear literature-supported function (Supplementary Figure 4C).

**Lymphocyte (187,237 cells):** the atlas contained five lymphoid subtypes: CD8+ cytotoxic T cells (CD8+ Tc), CD4+ helper T cells (CD4+ Th), CD4+ regulatory T cells (CD4+ Treg), naïve T cells and plasma cells. Atlas transfer fell below threshold (overall recovery 0.62), so the compartment was annotated *de novo*. Clusters were annotated as T.CD8 (*CD8A*+, *GZMK*+, and *CTSW*+), T.CD4 (*CD40LG*+, *IL32*+, and *IL2RA*+), T.NK (*CTSW*+, *KLRB1*+, *ID2*+, and *FCER1G*+, with low or negative *CD8A* and *CD3E*), B.Lymphocyte (*CD79A*+, *IRF8*+, and *MZB1*+), T.Naive (*CCR7*+, *LTB*+), T.Cycling (*MKI67*+, *TOP2A*+, *STMN1*+) and T.NOS (*CD3E*+). Cluster 13 was annotated as fibroblasts (*COL6A1*+, *PDGFRA*+, and *CCL19*+ with neglectable levels of *CD3E*) (Supplementary Figure 4D).

**Myeloid (169,435 cells):** the atlas resolved macrophages, monocytes, Langerhans cells (LC), and three dendritic-cell (DC) populations (*CD1C*+ DC, *CLEC9A*+ DC, *CCR7*+ DC), transferred at an overall recovery of 0.83. Monocytes were merged into macrophages (Mac) as the Xenium panel does not contain monocyte-specific markers, and *CD1C*+ DC was renamed cDC2 as *CD1C* was not included in the panel. Upon further inspection, cluster 6 was overruled to B.Lymphocytes (*CD79A*+/*MZB1*+) and cluster 13 to plasmacytoid DC (pDC, *IL3RA*+/*IRF8*+) (Supplementary Figure 4E).

**Epithelial (306,164 cells):** the atlas included thirteen subtypes: basal keratinocytes (KC), spinous (Spn) KC types I-II, granular (Grn) KC, cycling (cyc) KC, basal infundibulum (Inf), differentiated Inf, basal and suprabasal outer root sheath (ORS), inner root sheath (IRS), bulge, eccrine (Ecc) ducts and Ecc glands, transferred at an overall recovery of 0.72. After coarsening mixed components (Spn KC I/II and Grn KC into Suprabasal KC, basal Inf and differentiated Inf into Inf, the two ORS states into ORS), recovery rose to 0.86 and the transfer was accepted. Three clusters were overruled after transferring based on marker expression: cluster 16 was named Melanoma (*MLANA*+/*PMEL*+/*TYR*+), cluster 22 was overruled to Cyc KC (*MKI67*+/*NUSAP1*+/*TOP2A*+), and cluster 13 was overruled to Suprabasal KC (*KRTDAP*+/*DSC1*+) (Supplementary Figure 4F).

##### Melanoma gene co-detection metaprogram analysis

To identify transcriptional metaprograms within melanoma cells, we constructed a gene co-detection network, based on binary gene-detection patterns (Qiu, 2020) but using different edge weights. For each pair of genes, we computed Cohen’s kappa statistic on binarized detection (a gene is called detected in a cell when its count number exceeds zero), measuring agreement of two genes being jointly detected or jointly non-detected beyond what is expected from the marginal detection frequency of each gene (chance) (Cohen, J., 1960). The resulting gene–gene kappa matrix was used to construct a weighted, undirected co-detection network. Edges were retained for gene pairs with kappa values above an empirical threshold (threshold = 0.15, corresponding to the mean plus 2.3 standard deviations of the off-diagonal kappa values) (Supplementary Figure 5A). The largest connected component of this network was retained, and genes with one or fewer connections were excluded from community detection to focus on recurrent multi-gene structure rather than isolated or weakly connected genes (122 of 260 genes). Weighted Louvain community detection (Blondel et al., 2008) was then applied to the weighted network which partitioned the remaining 122 genes into five initial modules (Supplementary Figure 5B).

Module cohesion was assessed by conductance (Yang and Leskovec, 2012), calculated as:

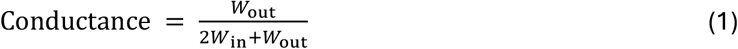

where *W*_in_ and *W*_out_ denote the summed weights of internal and external edges, respectively. Lower values indicate stronger separation from the rest of the network. Four modules were internally connected (conductance < 0.25), whereas module 3 was markedly less cohesive (conductance = 0.44), motivating further examination of its internal structure (Supplementary Figure 5C). Therefore, module 3 was re-clustered following the same approach (kappa matrix, k > 0.15 threshold, Louvain clustering), dividing it into three submodules (Supplementary Figure 5D) and yielding seven melanoma metaprograms in total (Fig. 2A–C). Metaprogram scores were calculated from log-normalized expression using Seurat’s AddModuleScore function (ctrl = 5, nbin = 10). Each melanoma cell was assigned to the metaprogram with the highest score.

##### Annotation of melanoma metaprograms

To assign biological identities to the seven malignant metaprograms, we tested each metaprogram for gene membership overlap against publicly available melanoma signatures. To that end, we combined ten single-cell and bulk transcriptomic studies that together span the major reported axes of melanoma phenotypic heterogeneity (Hoek, K.S., et al., 2008), (Verfaillie, A., et al., 2015), (Tirosh, I., et al., 2016), (Tsoi, J., et al., 2018), (Jerby-Arnon, L., et al., 2018), (Rambow, F., et al., 2018), (Wouters, J., et al., 2020), (Belote, R.L., et al., 2021), (Karras, P., et al., 2022), and (Pozniak, J., et al., 2024). Studies reporting gene identifiers other than human symbols were mapped during harmonization: the mouse gene sets of Karras et al. were converted to human orthologs, and Ensembl gene identifiers from Rambow et al. were resolved to HGNC symbols. This yielded 58 melanoma-state signatures. This reference set was expanded with the 41 Gavish pan-cancer metaprograms (Gavish, A., et al., 2023) and the 50 MSigDB Hallmark gene sets (Liberzon, A., et al., 2015) to assess broader transcriptional programs and pathways. Gene sets were obtained directly from the publications and the MSigDB database (msigdbr R package). Because the overlap can only be evaluated against the 260-gene Xenium panel, all overlap statistics used that panel as the background set rather than the whole transcriptome. All 149 reference signatures were retained in the direct metaprogram-overlap tests, including those with no genes represented in the panel. Pairwise comparisons were restricted to the 136 reference signatures with at least one represented gene, together with the seven metaprograms. Signature sizes, panel coverage, overlap counts and complete metaprogram–reference overlap results are provided in Supplementary Table 3. For each public signature, we counted the number of genes present in the Xenium panel (N) and for each metaprogram-signature pair we counted the shared panel genes (X). We used these two values to quantify the Dice coefficient as:

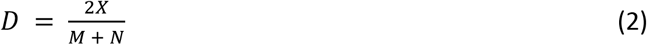

where M is the number of genes in the metaprogram. Statistical significance was assessed with a one-sided hypergeometric test, and p-values were corrected across all 1,043 metaprogram–reference pairs using the Benjamini-Hochberg (BH) method (Fig. 2D; Supplementary Table 3). To visualize the relationship among public signatures and metaprograms directly, we additionally computed pairwise Dice similarities and one-sided hypergeometric enrichment p-values across all 10,153 unordered pairs of represented signatures and metaprograms, with a separate BH correction across this complete family (Supplementary Figure 5E) and rendered these as a force-directed similarity graph, retaining edges with adjusted p-value < 0.1 (Supplementary Figure 5F).

Further, we projected the 7 Xenium melanoma metaprograms onto the Kunz nevus-melanoma cohort (Kunz, M., et al., 2018), comprising 57 primary melanoma and 23 melanocytic nevus profiles. Processed gene expression data and sample annotations were downloaded from GEO (GSE112509). Expression values were log1p transformed and mapped to gene symbols to match the metaprogram definitions. Metaprogram scores were obtained for each sample using AddModuleScore in Seurat which were subsequently scaled (z-score) and trimmed to the [-3, +3] interval to prevent a strong impact of any potential outlier in downstream analyses. Each sample was assigned a dominant metaprogram corresponding to the one with the highest scaled score (Supplementary Figure 5I). We compared melanomas and nevi using both dominant-metaprogram assignments and continuous metaprogram scores. For each metaprogram, the proportion of expression profiles assigned to that metaprogram was compared between melanomas and nevi using a two-sided Fisher’s exact test across the 57 melanoma and 23 nevus profiles (Supplementary Fig. 5I). Separately, continuous scores were averaged within each patient and lesion class, yielding 51 melanoma and 23 nevus observations. These were compared using a linear model with lesion class as the predictor, equivalent to an unpaired, equal-variance Student’s t-test; effect sizes were expressed as nevus minus melanoma (Supplementary Fig. 5H; Supplementary Table 3). For each analysis, P values were adjusted across the seven metaprograms using the Benjamini–Hochberg method.

Metaprograms were annotated by interpreting their constituent gene sets, overlap with public melanoma transcriptional signatures, spatial localization, and projection onto the independent GSE112509 nevus– melanoma cohort. The resulting annotations were: module 1, differentiated melanocytic; module 2, epithelial-adjacent; module 3, mesenchymal-invasive; module 4, proliferation/cell cycle; module 5, early developmental stress; module 6, nevus-like; and module 7, IFN-associated stress.

##### Metaprogram trajectories

Next, we assessed the location of the melanoma metaprograms based on their distance to the epidermis. To that end, we calculated the median Euclidean distance of each melanoma cell to its closest ten suprabasal keratinocytes. This comparison was limited to the first 400µm of depth which already included 90.6% of melanoma cells and excludes a long, low-density tail (Supplementary Figure 7A). Metaprogram composition was evaluated as the percentage assigned to each metaprogram within each 25-µm bin. Bins containing fewer than 250 melanoma cells were excluded. Depth-dependent proportions were visualized using loess smoothing with a span of 0.45.

Given the different spatial locations of melanoma states, we examined whether they could be ordered along inferred transcriptional trajectories. To that end we applied Slingshot (Street, K., et al., 2018) on a 3-dimensional diffusion map (DM, destiny R package) (Angerer, P., et al., 2016) built on the normalized metaprogram scores (z-score, capped ±3) from a subset of up to 1,000 randomly selected cells per metaprogram. The initial diffusion map separated the proliferation/cell-cycle program from the other metaprograms (Supplementary Figure 7B). This is in line with previous reports showing that cell-cycle-associated variation can obscure differentiation-related structure and affect trajectory inference (Charrout et al., 2020). We therefore repeated the diffusion map and Slingshot analyses using the six remaining metaprogram scores, excluding cells assigned to the proliferation/cell-cycle metaprogram from model fitting and using epithelial-adjacent as the root cluster. Slingshot returned two lineages sharing a common trunk (Supplementary Figure 7C). The shared trunk included: epithelial-adjacent > early developmental stress > differentiated melanocytic, which then split into: IFN-associated stress > mesenchymal-invasive (lineage 1) (Supplementary Figure 7D); and nevus-like (lineage 2) (Supplementary Figure 7E). The fitted model was projected onto the complete melanoma dataset where each cell was assigned a pseudotime on each lineage (t_1_, t_2_) and a curve weight (w_1_, w_2_). From these, we defined a merged progression axis and a branch axis:

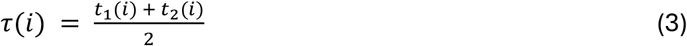

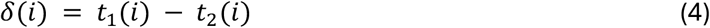

Because the two lineages share the trunk, each trunk cell carries two pseudotime estimates that should agree there (t1≈t2, i.e. δ≈0). In practice t2 ran systematically ahead of t1, an offset between the inferred pseudotime coordinates that increased along the trunk (Supplementary Figure 7F). This was addressed by aligning the two trajectories along the shared trunk. To that end, we first defined trunk cells as those having joint curve weights (w1+w2) >= 1.5. We ordered trunk cells along the progression axis (τ) and grouped them into 70 equal-frequency bins; within each bin we took the median t1 and the median t2, retained bins containing at least 20 cells, and smoothed the resulting sequence of bin medians (loess, span = 0.25), tracing a single curve S through the trunk cells in the plane. For each cell, we identified the nearest point on the piecewise-linear trunk skeleton S, with coordinates s1(i) and s2(i). The local coordinate difference at that point was subtracted from t2:

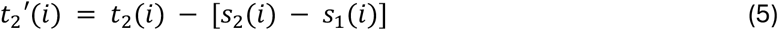

This aligned the shared trunk with the diagonal (Supplementary Figure 7G). The progression and branch axes were then recalculated using the corrected t2 values for fork detection and trajectory classification.

To identify the fork where both lineages split, we calculated the dispersion of the branch axis (*δ)* along the progression axis (*τ).* Cells were binned into 100 quantile bins of τ and per bin dispersion was computed using the median absolute deviation (MAD). The MAD curve was loess-smoothed (span = 0.3) for visualization. The fork was estimated as the first bin-median progression value where the unsmoothed MAD exceeded 10% of its maximum (Supplementary Figure 7I), which returned a value of τ_fork = 0.056. Each cell was assigned a trajectory class (pt_class) from its progression and curve weights:

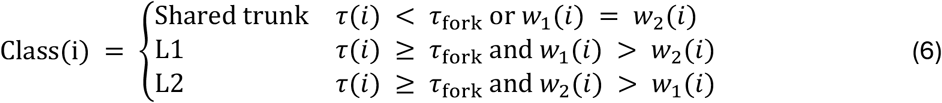

Cells unusually far from their initially assigned trajectory skeleton were reassigned and their pseudotime coordinates adjusted. We constructed smoothed skeletons for the shared trunk (70 bins; loess span = 0.25) and each lineage (90 bins; span = 0.30), and measured the Euclidean distance from each cell to the nearest vertex of its assigned skeleton. Cells exceeding the pooled mean distance plus three standard deviations (threshold = 0.0081; 1,001 cells, 0.45%) were reassigned to the lineage with the larger corrected pseudotime coordinate, with ties assigned to Lineage 1. Both coordinates were replaced by those of the skeleton vertex closest along that dominant coordinate (Supplementary Figure 7H–J).

The final pt_class assignments (Shared Trunk 98,700; Lineage1 64,433; Lineage2 57,786) were cross-tabulated against the metaprogram labels. Enrichment or depletion was quantified as the observed-to-expected ratio under independence (O/E). The overall association was assessed using a χ² test, with Cramér’s V reported as a global effect-size measure.

##### Cross-run reproducibility

After second-level clustering and melanoma metaprogram characterization, cells were resolved into 42 cell-type annotations across eight broad lineages (Supplementary Fig. 8D). All 10 patients were acquired in two or more independent Xenium runs (21 acquisitions in total): nine were acquired twice and one (thinmme008) was acquired three times. Pairwise run-vs-run comparisons were performed within each patient, giving 12 matched run-pairs. Per-run-pair concordance was computed from the level-2 cell-type composition of each acquisition (percentage of cells assigned to each label) using the Aitchison distance (Aitchison, J., 1994) which calculates the Euclidean distance between centered log-ratio (CLR) transforms of the cellular proportions (a residual of 1×10⁻⁶ added and re-normalized before the CLR to avoid taking the logarithm of zero), which is scale-invariant and appropriate for compositional data:

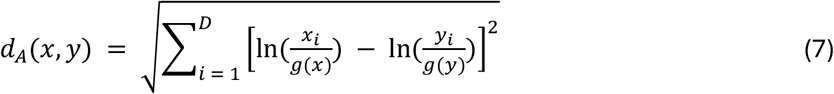

where D is the number of cell-type annotations, g(x) is the geometric mean of composition x, and 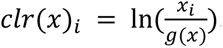.

To test whether replicate acquisitions of the same patient were compositionally more similar to each other than to acquisitions from different patients, we compared the Aitchison distance between matched runs and mismatched runs (acquisitions from different patients) and derived an empirical p-value over 100,000 permutations of the patient labels, counting how often the permuted mean matched-run distance was ≤ the observed value (mean matched 3.30 vs. mismatched 7.68; p < 1×10⁻⁴) (Supplementary Fig. 8E). Despite the overall concordance, run-to-run agreement was patient-dependent. Most samples reproduced closely (e.g. thinmme006, mean Aitchison distance 2.40) (Supplementary Fig. 8G), whereas thinmme001 showed the largest run-to-run discordance (Aitchison distance 8.50) (Supplementary Fig. 8H), consistent with reduced tumor representation in its second acquisition (the tumor was almost exhausted in run 2). To avoid propagating this variability into downstream composition analyses, we excluded acquisitions having less than 1,000 melanoma cells and fewer than 25% of the maximum melanoma cell count in the samples of that patient. This removed two acquisitions (thinmme005 run 1 and thinmme001 run 2), leaving 19. Cell counts were then pooled across retained acquisitions within each patient, and subtype proportions were calculated within each broad cellular compartment.

##### Paired compositional differential abundance analysis

Compositional differences between M+ and M-tumors were tested with propeller (Phipson, B., et al., 2022), which models arcsin-square-root-transformed cell-type proportions per sample within a moderated-t (limma empirical-Bayes) framework. Because the study design pairs each M+ patient with a matched M-patient, we adapted the propeller implementation in the speckle R package enabling paired comparisons (∼ met + pair) rather than using a simple two-group comparison. Testing was performed within each of the eight level 1 lineages separately, plus a parallel melanoma pseudotime branch classification. P-values were corrected for multiple testing using the Benjamini–Hochberg (BH) method separately within each level 1 lineage and within the melanoma pseudotime branch classification.

##### Spatial neighborhood analysis

###### Spatial neighborhood enrichment

For each melanoma cell in each retained acquisition, we identified the 25 nearest other cells using Euclidean distances between cell centroids. Neighbors could belong to any of the 42 cell annotations. We grouped melanoma reference cells by metaprogram and compared their neighbors with the cell composition of the same acquisition. Enrichment was calculated as:

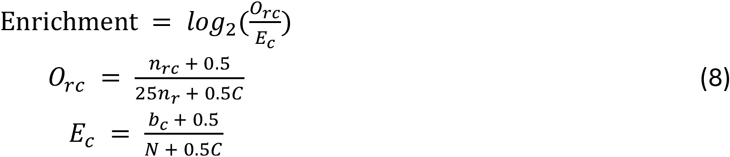

where *O_rc_* is the observed neighbor frequency and *E_c_* is the expected frequency from the same acquisition. Here, *n_rc_* is the number of neighbors annotated as c around reference cells in group r, *n_r_* is the number of reference cells in that group, *b_c_* is the number of cells annotated as c in the acquisition, N is its total cell count, and C = 42. Adding 0.5 to each category before normalization avoids zero frequencies. We summarized enrichment as the median across acquisitions; displayed neighbor counts are the median across acquisitions of the mean count per reference cell (Supplementary Fig. 8A). We repeated the analysis with reference cells grouped as shared trunk, lineage 1 or lineage 2, keeping the same 42 neighbor annotations (Supplementary Fig. 8B).

###### Continuous-pseudotime neighborhood evolution

We used all retained melanoma cells to describe how neighborhood composition changed along each trajectory. Shared-trunk cells contributed to both trajectories, while branch-assigned cells contributed only to their corresponding lineage. Using the respective lineage-specific pseudotime coordinates, we grouped reference cells into bins of 0.002 pseudotime units and pooled counts across acquisitions within each patient. Patient–lineage–bin combinations with fewer than ten reference cells were excluded. For each neighbor type, we calculated its percentage among the 25 neighbors per reference cell and took the median across all patients meeting the reference-cell threshold, including those with zero neighbors of that type. Profiles were displayed with quadratic loess smoothing (span = 0.75) (Fig. 2I). We compared the unsmoothed bin-level percentages between lineages using:

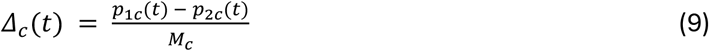

where p₁c(t) and p₂c(t) are the median neighbor percentages for type c in pseudotime bin t along lineages 1 and 2. Mc is the highest percentage for that neighbor type across both lineages and all displayed bins. Thus, positive values indicate a higher percentage along lineage 1 and negative values a higher percentage along lineage 2 (Supplementary Fig. 8C). Only bins supported by both trajectories were compared. If a neighbor type had zero median percentages in both trajectories across all displayed bins, its normalized difference was set to zero.

#### Milan Analysis

Tissue sections were stained using a 42-antibody MILAN panel encompassing cell-lineage and cell-state markers (Supplementary Table 9).

##### Image preprocessing

Image analysis was performed using the custom DISSCOvery pipeline (Antoranz, A., et al., 2022). Briefly, flat-field correction (FFC) was performed on raw fluorescence tiles using BaSiC (Peng, T., et al., 2017). Tissue foreground was segmented from background using a custom model, and slides having multiple tissue sections were divided into separate tissue regions for downstream steps. The final acquisition round (R17) was used as a reference to define tissue bounding boxes, foreground masks, and registration fixed images. Imaging artifacts, including air bubbles, antibody aggregates, tissue folds, external artifacts, and out-of-focus regions, were identified using QUALIFAI (Andhari, M., et al., 2024) and excluded from downstream analysis. Per-sample tiles were stitched and registered across cycles using COLLAGE (Antoranz, A., et al., 2024), with DAPI as the alignment channel. The resulting transformation matrices were applied to the remaining channels. Affine alignment was subsequently refined by non-rigid registration using VALIS (Gatenbee, C.D., et al., 2023). Segmentation was performed using a fine-tuned StarDist v0.7.3 model (Schmidt, U., et al., 2018). Autofluorescence was subtracted using CLEAR-AF (Arnould, A., et al., 2025), with baseline images selected according to acquisition proximity and adjusted where necessary following visual quality assessment. Finally, spatial coordinates (centroid X/Y), morphological features (segmented-object area), and molecular features (mean fluorescence intensity of each marker within the segmented object, MFI) were extracted.

Staining quality was assessed by visual inspection of individual marker images and multiplex overlays, considering signal intensity, background and localization within the tissue architecture (Fig. 3C). Nine markers failed QC and eight had uncertain staining quality; neither group was used in downstream analysis. The remaining 25 markers passed QC (Supplementary Table 9).

##### Cell Phenotyping

Cells with zero MFI across all measured markers were excluded before cell phenotyping. MFI values were normalized separately for each marker within each tissue region to z-scores and clipped to the range [−5, 5] to limit the influence of extreme values (Caicedo, J.C., et al., 2017). A subset of 25,000 cells was selected by proportional stratified random sampling across tissue regions. Cell identification used 17 phenotypic markers: AQP1, αSMA, CD1a, CD3, CD4, CD20, CD31, CD68, CD79a, CK, FAP, FOXP3, MELANA, MITF, S100, S100B and SOX10.

Cells were clustered using CONCLAVE (Nazari, P., et al., 2026), a consensus approach combining three complementary clustering methods: PhenoGraph, K-means and SOM-based clustering. Before annotation, cluster labels from each method were projected to the full dataset using nearest-neighbor voting in a UMAP embedding fitted to the sampled marker-expression profiles (nneighbors = 25). Projected clusters were visualized at their original tissue coordinates to assess their spatial distribution.

Clusters were manually assigned to cell types by jointly evaluating their marker-expression profiles and localization within the tissue architecture. This identified tumor cells (MELANA+, MITF+, S100+, S100B+ and/or SOX10+), epithelial cells (CK+), endothelial cells (CD31+ and/or AQP1+), fibroblasts (FAP+), smooth-muscle cells (αSMA+), macrophages (CD68+), B cells (CD20+ and/or CD79a+), helper T cells (CD3+ and CD4+), cytotoxic T cells (CD3+ and CD4-, CD8 did not pass QC) and regulatory T cells (CD3+, CD4+, and FOXP3+), together with Blank and Noise categories. Cells assigned the same broad phenotype by at least two of the three clustering methods were retained as a consensus reference.

For final cell-type assignment, the consensus reference was subsampled to a maximum of 2,500 cells per phenotype and embedded in a three-dimensional UMAP space. Reference labels were refined using agreement among 25 nearest neighbors. The full dataset was subsequently projected into this reference embedding, and phenotypes were assigned by nearest-neighbor voting (nneighbors = 25), with prediction confidence recorded for each cell. Patient-level MILAN counts and proportions are provided in Supplementary Table 5.

#### Cross-modal concordance and integration of MILAN protein and Xenium RNA profiles

Independently derived MILAN and Xenium annotations were mapped to shared lineage categories and compared across 217,798 spatially matched cell pairs. Concordance was calculated as the percentage of cell pairs assigned to the same lineage by both modalities, overall and for each Xenium lineage. We calculated agreement both including all pairs and after excluding MILAN Blank and Noise assignments. Overall agreement was 53.9% across all pairs and 78.6% among the 149,222 pairs with classified MILAN annotations.

RNA–protein correspondence was assessed for five directly matched marker pairs: *MLANA*/MELANA, MITF, S100B, AQP1 and *MKI67*/Ki67. Pan-cytokeratin staining was additionally compared with the mean normalized Xenium expression of *KRT5*, *KRT15* and *KRT18*. Pearson correlations were calculated between normalized Xenium RNA expression and normalized MILAN protein measurements across matched cells, both pooled across patients and separately within each patient. Pooled correlations ranged from 0.47 to 0.69, while patient-specific correlations ranged from 0.37 to 0.81 (Supplementary Fig. 9G–J).

To compare the spatial distribution of *MLANA* RNA and MELANA protein in thinmme003, we used raw transcript counts and fluorescence intensities from 22,297 matched Xenium-defined melanoma cells. Each measurement was divided by its modality-specific mean across these same cells. Relative RNA versus protein enrichment was then calculated as:

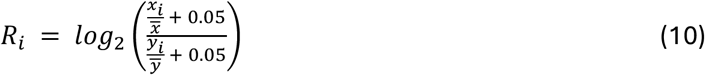

where *x_i_* and *y_i_* denote the *MLANA* transcript count and MELANA fluorescence intensity of matched cell *i*, respectively, and *x* and *y* are their means across the matched melanoma cells. The residual of 0.05 prevented divisions by 0 or taking the logarithm of 0. Positive values indicate greater relative RNA enrichment and negative values greater relative protein enrichment.

Protein profiles of Xenium-defined melanoma metaprograms were evaluated using MELANA, MITF, S100, S100B, SOX10, SOX9, AQP1, pan-cytokeratin, Ki67, NGFR and TCF4. Cells were grouped by Xenium melanoma metaprogram. Protein measurements were z-scored separately for each marker across matched melanoma cells, and mean z-score was calculated within each metaprogram (Fig. 3H).

For trajectory comparisons, RNA and protein measurements were z-scored separately for each marker and modality across matched melanoma cells. Cells were then grouped by trajectory assignment, with shared-trunk cells contributing to both lineages. Merged pseudotime was rounded to 0.001-unit bins, and mean z-score RNA and protein were calculated within each lineage and bin (Fig. 3J).

#### VISIUM analysis

##### Quality Control and normalization

Visium CytAssist data were generated from serial sections corresponding to three matched pairs of M+ and M-thin melanomas. Visual inspection of the H&E images confirmed that the captured spots were located on tissue. However, the sections included substantial areas of naturally hypocellular deep dermis and residual adipose tissue. These regions produced lower numbers of detected genes and transcripts despite representing genuine tissue (Bhuva et al., 2024) (Supplementary Figure 12A-B). To focus the analysis on the epidermal, tumor and tumor-associated microenvironmental compartments, only spots with at least 750 detected genes were preserved (Supplementary Figure 12C). The final focused dataset contained 7,652 spots across the six sections (out of 22,749 on tissue spots; fraction retained = 33.64%).

Raw counts were normalized independently for each section using SCTransform v2 with the glmGamPoi implementation (Hafemeister and Satija, 2019), (Ahlmann-Eltze and Huber, 2020). A shared set of 3,000 genes was selected using cross-section variable-feature rankings, limiting the selection to genes detected in at least five spots in every section. Pearson residuals for these genes were retained for dimensionality reduction and integration.

##### Integration and Dropout Imputation

Despite section-wise normalization, the unintegrated SCT representation retained substantial section-specific structure (Supplementary Figure 12D). We therefore integrated the six sections using reciprocal principal-component analysis (RPCA) on the SCT representations (Stuart et al., 2019). Integration was performed using 3,000 shared residual features and 30 principal components, identifying 15,512 anchors across sections. Local cross-section mixing was quantified using kBET in the first five principal components, with 30-spot neighborhoods and 100 repetitions (Büttner et al., 2019).

To reduce the influence of expression sparsity (Wang et al., 2022), MAGIC (van Dijk et al., 2018) was applied to section-wise SCT expression using a joint neighborhood graph built on the 30-dimensional RPCA embedding. This produced imputed expression for 18,085 genes shared across all six sections in 7,652 QC-passing spots. Imputation was evaluated against registered Xenium data from the paired sections. Using the cell-to-spot assignments described above (see Cross-modal alignment), raw counts were summed across Xenium cells assigned to each Visium spot, followed by library-size normalization and log1p transformation. Agreement between Visium and Xenium pseudospot expression across 252 shared genes was assessed using Lee’s L, a spatially weighted bivariate statistic that measures correspondence between local expression patterns (Lee, 2001).

##### Xenium-guided Lineage Deconvolution

Because Visium spots contain mixtures of multiple cell types, broad lineage composition was estimated using the matched Xenium dataset as a single-cell reference. The level 1 Xenium annotations were used which include 7 categories: melanoma, epithelial, fibroblast, lymphocyte, myeloid, endothelial, and other stromal cells. From each L1 lineage and patient, up to 1,000 cells were sampled at random to prevent abundant populations or individual specimens from dominating the reference. Lineage abundances were inferred from raw Visium counts using cell2location (c2l) (Kleshchevnikov et al., 2022), and the resulting values were normalized within each spot.

To identify melanoma-associated spots, the continuous c2l melanoma weight was calibrated against the registered Xenium cells. Spots containing at least five registered melanoma cells, representing at least 25% of their registered cells, defined the reference tumor ground truth. Threshold selection for the c2l melanoma weights was based both on F1 score and the Matthews correlation coefficient (MCC), which is a more robust metric under class imbalance than F1 alone (Chicco and Jurman, 2020). Both metrics peaked at low thresholds (F1 maximal at 0.19, F1 = 0.555; MCC maximal at 0.14, MCC = 0.464); a relative melanoma-weight of 0.20 was nonetheless selected as a stringent cutoff above both optima (F1 = 0.553, MCC = 0.458 at this threshold), trading some sensitivity for a purer tumor-domain definition. This threshold identified 1,590 melanoma-associated spots and retained 70.2% of the reference tumor-positive spots in the registered subset (Supplementary Figure 13B). These spots defined the melanoma-focused domain used for subsequent metaprogram analyses.

##### Melanoma Metaprogram Analysis

The seven melanoma metaprograms identified in the matched Xenium dataset were projected onto the 1,590 melanoma-associated Visium spots using their gene signatures. Metaprogram scores were calculated on the MAGIC-imputed expression using AddModuleScore from Seurat. Because the score distributions differed between metaprograms and sections, the Visium scores were calibrated separately for each patient and metaprogram. Visium score ranks were mapped onto the corresponding score distribution measured across all Xenium melanoma cells from the same tumor. Spots were assigned to a metaprogram when their calibrated score exceeded a threshold, selected to maximize agreement between Xenium metaprogram compositions, assessed using Spearman correlation and mean absolute error (MAE) (threshold = 0.15; Supplementary Figure 13C–D). Since each metaprogram was assigned independently, individual spots could be assigned to more than one metaprogram, in line with the multicellular nature of Visium spots. Of the 1,590 melanoma-associated spots, 554 (34.8%) had at least two memberships, including 131 (8.2%) with at least three and 30 (1.9%) with four; none had more than four. A further 676 spots (42.5%) were assigned to a single metaprogram, while 360 (22.6%) did not exceed the assignment threshold for any metaprogram. Patient-level Visium metaprogram composition was calculated as the fraction of melanoma-associated spots assigned to each metaprogram and compared with the corresponding fractions among Xenium melanoma cells.

The Visium spots were subsequently projected onto the fixed Xenium-derived diffusion map and Slingshot trajectory. Visium spots were subjected to the same corrections performed for the Xenium cells (see corresponding section in Xenium analysis). Spots were classified as shared trunk or as one of the two lineages using the same pseudotime threshold established on the Xenium trajectory, classifying 661 spots along the shared trunk, 358 along lineage 1, and 571 along lineage 2 (Fig. 4B; Supplementary Figure 13E). Patient-level lineage composition was calculated as the fraction of Visium spots assigned to each of the three trajectory classes and compared with the corresponding fractions of Xenium melanoma cells.

##### Functional Characterization of Melanoma Metaprograms and Trajectories

To characterize the broader transcriptional context of the Xenium-defined melanoma metaprograms, pathway activity was evaluated across the 1,590 melanoma-associated Visium spots using pre-MAGIC, section-wise SCT expression. GSVA scores were calculated for the 50 MSigDB Hallmark gene sets (Hänzelmann et al., 2013; Liberzon et al., 2015). For each metaprogram, assigned spots were compared with the remaining melanoma-associated spots using two-sided Wilcoxon rank-sum tests, with Benjamini– Hochberg correction across pathways within each comparison. Effect sizes were expressed as differences in median GSVA scores (Fig. 4C). Gene-level member-versus-rest comparisons were also performed on pre-MAGIC SCT expression using Wilcoxon rank-sum tests, with Benjamini–Hochberg correction across 18,085 genes within each comparison. Complete results are provided in Supplementary Table 6.

Gene-expression changes along the two melanoma trajectories were assessed using tradeSeq negative-binomial generalized additive models fitted to raw counts and corrected lineage pseudotimes (Van den Berge et al., 2020). Genes detected in at least 10% of the 1,590 melanoma-associated spots were modeled, retaining 11,980 genes. Shared-trunk spots were assigned to both lineages. Chi-square upper-tail probabilities were calculated directly, and P-values were adjusted using the Benjamini–Hochberg procedure separately within the global, Lineage 1, Lineage 2 evaluations. Complete results are reported in Supplementary Table 6. Gene curves show expected counts at the median observed library-size offset, evaluated over the 2.5th–97.5th percentiles of sampled pseudotime within each lineage. Hallmark GSVA scores were modeled using Gaussian GAMs fitted separately along each lineage with cubic shrinkage splines (basis dimension k = 5), estimated by restricted maximum likelihood. Shared-trunk spots contributed to both lineage datasets, giving 1,019 Lineage 1 and 1,232 Lineage 2 observations. Approximate smooth-term tests were followed by Benjamini–Hochberg correction across the 100 pathway–lineage tests. The fitted score range reported in Supplementary Table 6 was the maximum minus minimum predicted GSVA score over the 2.5th–97.5th percentile pseudotime interval within each lineage, capturing both peaks and troughs. The four pathways displayed in Figure 4F were selected to illustrate shared trends (MYC targets V1 and EMT) and divergent profiles (inflammatory response and complement). All 100 pathway–lineage tests, including nonsignificant results, are reported in Supplementary Table 6. These tests assessed within-lineage associations and did not directly test pathway differences between lineages. The pooled spot-level models did not adjust for patient effects or spatial dependence.

#### International Multicentric Independent Validation

##### H&E-based inference of melanoma metaprograms

To explore whether the melanoma states identified by spatial transcriptomics could also be detected in H&E images, we applied DeepSpot-M (Nonchev et al., 2026) to an international multicentric melanoma cohort collected at UZ Leuven (Leuven, Belgium), the Maria Skłodowska-Curie National Research Institute of Oncology (Warsaw, Poland), and Careggi University Hospital (Florence, Italy). The outcome-labeled dataset comprised 60 slides representing 58 patients (35 M+ and 23 M−). DeepSpot-M predictions covered 257 decoder-supported genes from the 260-gene Xenium panel and were analyzed on the model’s output scale without further normalization.

To identify melanoma-enriched regions, we calculated the mean predicted expression of ten Xenium-derived melanoma markers: *TYR*, *MLANA*, *PMEL*, *NSG1*, *EDNRB*, *MITF*, *TYRP1*, *QPCT*, *PCDH7* and *TFAP2A*. The threshold was calibrated on the six Visium sections where we compared DeepSpot-M-based tumoral scores to the cell2location melanoma relative weights (spots were defined as melanoma enriched when weight ≥ 0.20). The threshold was selected maximizing Youden’s index (sensitivity + specificity − 1) (Youden, 1950) (threshold = 0.90424). This threshold was applied to the slides of the validation cohort. Within these spots, we calculated melanoma metaprogram scores using AddModuleScore (nbin = 10, ctr = 5). The seven signatures included 120 represented genes out of 122 requested genes; *MIR205HG* and *ZNF667-AS1* were unavailable. Each spot was assigned exclusively to its highest-scoring metaprogram. Metaprogram spot counts were summed across slides for the same patients before calculating fractions. Fractions were compared between M+ and M-using two-sided Wilcoxon rank-sum tests, with Benjamini– Hochberg correction. ROC curves were calculated. Confidence intervals were calculated using 5,000 outcome-stratified bootstrap resamples of patients.

##### Association of the nevus-like metaprogram with survival in external cohorts

To investigate the clinical relevance of the Xenium-derived programs, we evaluated the nevus-like signature using AddModuleScore in 703 primary melanomas from the Leeds Melanoma Cohort (Nsengimana et al., 2018) and 51 primary melanomas from GSE98394 (Badal et al., 2017), excluding the 27 benign nevi in the latter dataset. No thickness, stage or signature-dominance restriction was applied. Overall-survival analyses included all patients with valid positive follow-up: 703 Leeds patients (233 deaths) and 44 GSE98394 patients (18 deaths); seven GSE98394 cases lacked eligible survival data. Scores were analyzed continuously using full available follow-up. The supplied Leeds microarray expression matrix had undergone quantile normalization, singular-value-decomposition adjustment and log transformation; no additional log transformation was applied before module scoring.

Nevus-like scores were analyzed as continuous predictors in Cox proportional-hazards models. Separate models evaluated the unadjusted association, stratified by broad tumor-stage groups (I–III), or adjusted for log₂-transformed Breslow thickness. Kaplan–Meier curves compared patients in the lowest within-cohort decile of nevus-like scores with the remaining patients, using log-rank tests. The 10th-percentile cutoff was selected since 10% of metastatic melanomas were initially diagnosed as early-stage disease (Ertekin et al., 2021). Patient scores and clinical metadata are provided in Supplementary Table 7.

##### Estimated cellular contributions to bulk metaprogram expression

To estimate the contributions of melanoma cells to MES and nevus-like signature expression estimated from bulk gene expression, we estimated major-lineage cell fractions (melanoma, epithelial, endothelial, fibroblast, lymphocyte, myeloid, nerves and stroma) of bulk transcriptomic samples using MuSiC (Wang et al., 2019) and the Xenium data as the single-cell reference. We combined these fractions with the average expression of signature genes in each major-lineage cell fraction. Signature expression was calculated as the mean across its constituent genes.

Because Leeds used microarrays, we harmonized its expression distributions to synthetic Xenium mixtures using gene-wise matching of log-expression means and standard deviations before estimating cell fractions. This harmonization was used only for cellular attribution; the original expression data were retained for survival scoring.

For each patient, each lineage’s expected signature expression was calculated by multiplying its estimated cell fraction by its reference signature expression. Estimates were summarized across 703 Leeds and 51 GSE98394 profiles.

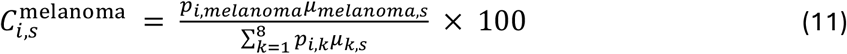

where 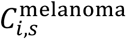 is the estimated percentage contribution of melanoma cells to signature s in patient i, *p_i_*_,*k*_ is the estimated cell fraction of lineage k, and *μ_k_*_,*s*_ is the mean Xenium reference expression of the genes belonging to signature s in that lineage.

## Data and code availability

Xenium, MILAN multiplex immunofluorescence and Visium data generated in this study will be deposited upon publication. We also provide an integrated object with all available modalities per sample in SpatialData format. Leeds Melanoma Cohort data are accessible through the European Genome-phenome Archive under accession EGAS00001002922.

Analysis scripts and code used to generate the insights of the study are available at https://github.com/antoranzlab/thin_melanoma

## Acknowledgements

We thank the patients whose tissue samples and clinical information made this study possible. The authors used ChatGPT (GPT-6.1 Sol) to assist with language editing. The study’s scientific concepts and design originated from the authors, who critically reviewed and edited all AI-assisted content and take full responsibility for the final manuscript, its interpretations and conclusions.

## Author Contributions

Conceptualization, A.A., M.D.A., and F.B.; methodology, A.A., M.D.A., and F.B.; software, A.A. and P.N.; validation, A.A., M.D.A., and F.B.; formal analysis, A.A., M.D.A., and P.N.; investigation (spatial-omics experiments, acquisition, and quality control), A.A., M.D.A., J.C., N.D., N.V., K.V., T.V., and F.B.; resources (validation samples and datasets), J.N., D.M., V.D.G., A.S.-C., P.R., P.T., A.M., and S.V.; data curation (patient identification and clinical metadata collection), A.A., M.D.A., C.G., P.G.D., D.V., A.L., J.P., D.M., V.D.G., A.S.-C., P.R., P.T., A.M., S.V., V.B., and F.B.; writing—original draft, A.A., M.D.A., and F.B.; writing—review and editing, all authors; visualization, A.A. and M.D.A.; supervision, A.A., O.G., and F.B.; project administration, A.A., M.D.A., and F.B.; funding acquisition, A.A. and F.B.

## Supplementary Information

**Supplementary Figure 1.**
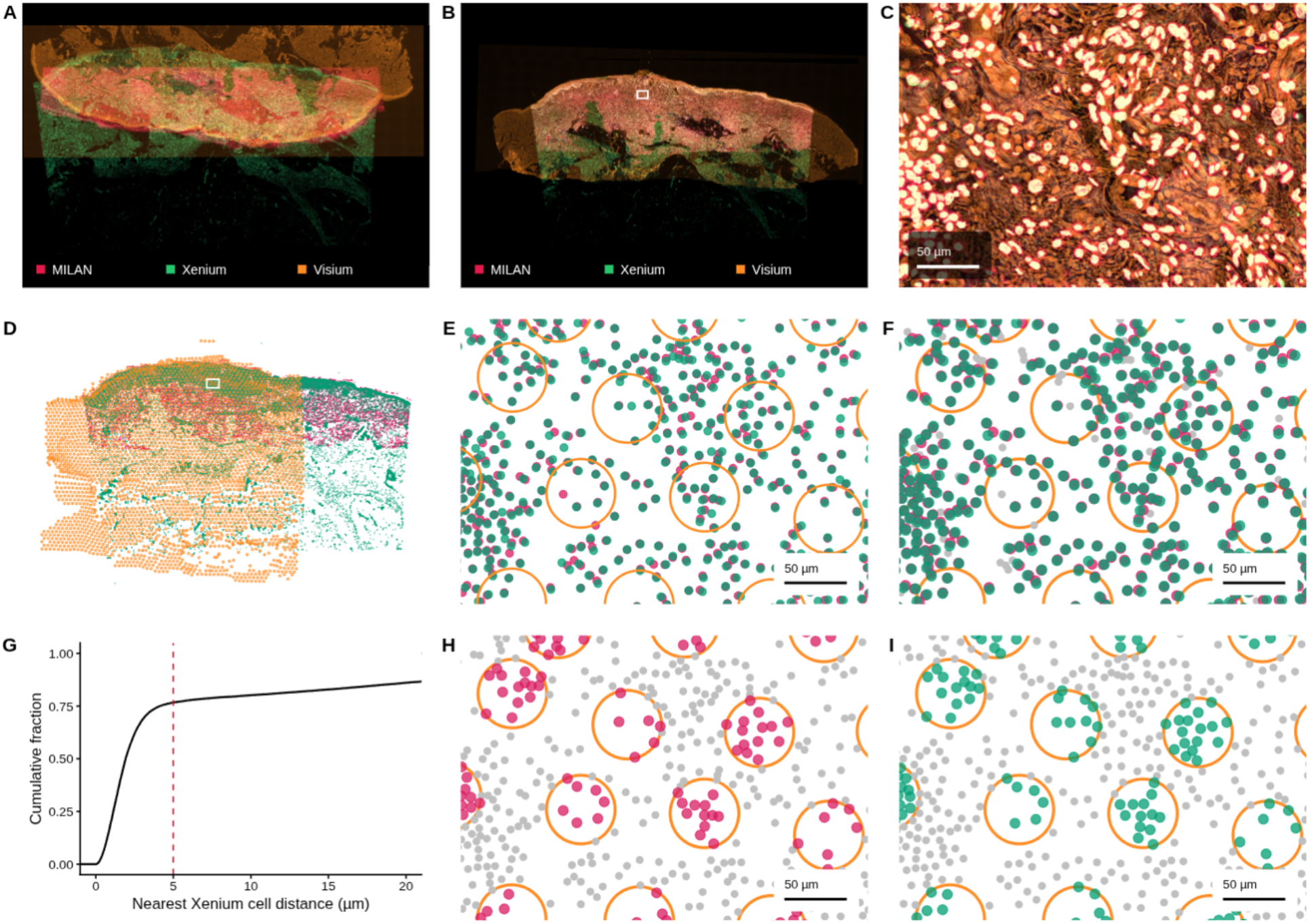
Cross-modal image registration and spatial assignment of Xenium, MILAN and Visium measurements. **A-B)** Overlay of MILAN and Xenium DAPI images and the Visium H&E image before (A) and after (B) registration. **C)** Magnified view of the region outlined in white in B. **D)** Registered MILAN and Xenium cell centroids overlaid with Visium spot footprints. **E)** Registered cell centroids and spot footprints within the selected region. **F)** Accepted Xenium– MILAN cell pairs, with matched cells colored by modality, unmatched cells in grey. **G)** Empirical cumulative distribution of the distance from each MILAN cell to its nearest Xenium cell across the illustrated sample, before applying the matching filters. The dashed vertical line marks the 5 µm distance threshold. **H–I)** Independent assignment of MILAN (H) and Xenium (I) cells to Visium spots. Colored cells have centroids within 27.5 µm of the spot center. Grey cells remain unassigned. Orange circles indicate Visium spots, with a diameter of 55 µm.

**Supplementary Figure 2.**
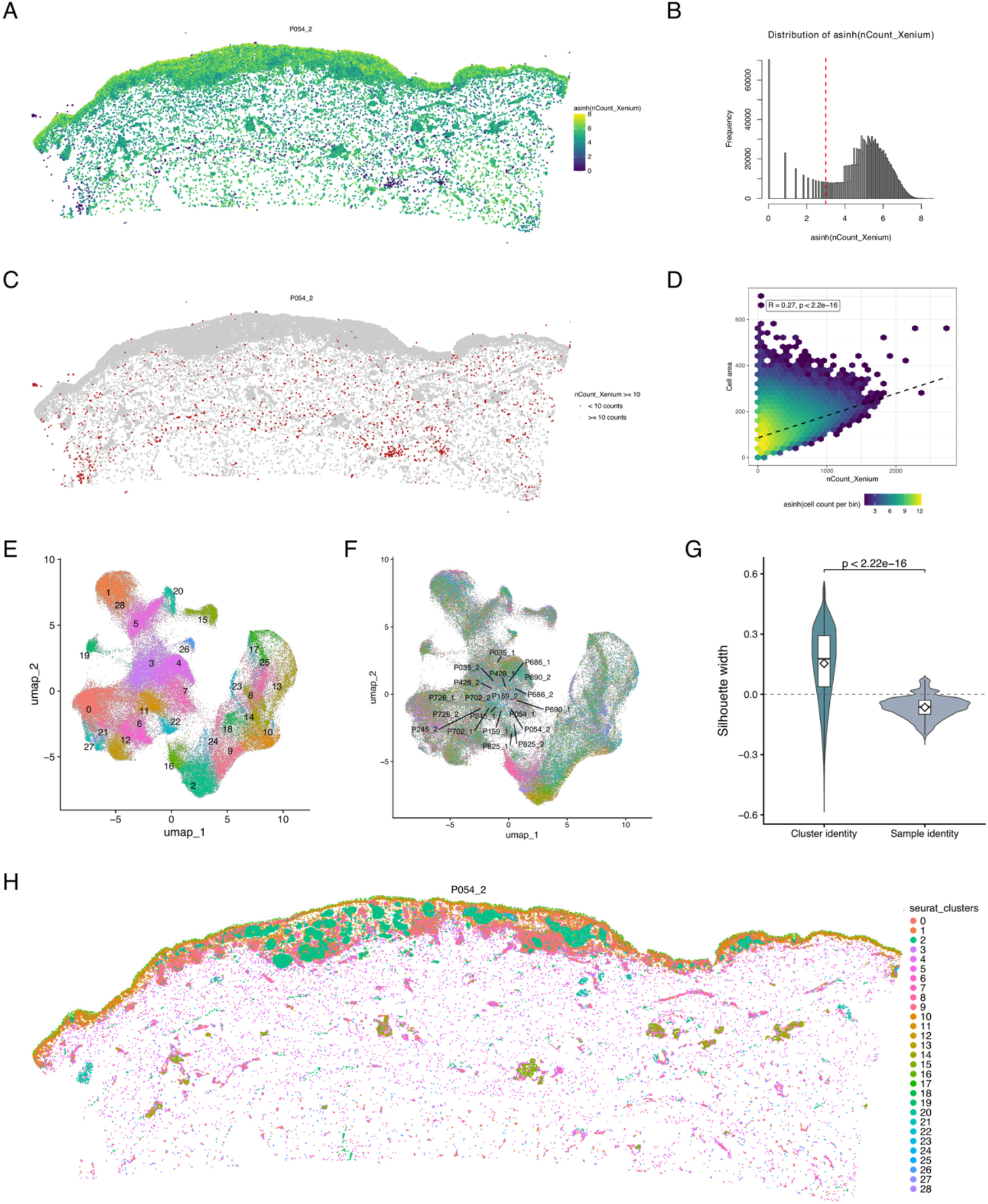
Quality control, normalization, and definition of first-level cellular phenotypes in Xenium dataset. **A)** Scatter plot showing the spatial distribution of total Xenium transcript counts per cell in sample thinmme006. **B)** Distribution of total Xenium counts per cell across the merged dataset. The red dashed line indicates the selected threshold (nCount_Xenium = 10). **C)** Scatter plot showing the spatial localization of cells passing the threshold in the same sample. **D)** Relationship between total Xenium counts and segmented cell area across cells. The dashed line indicates the fitted linear trend. **E)** UMAP embedding of a balanced reference subset colored by Seurat cluster. **F)** The same UMAP colored by sample identity. **G)** Quantification of the batch effect by Silhouette score. Cluster scores have significantly higher silhouette width in comparison with samples. **H)** Scatter plot of thinmme006 colored by cluster.

**Supplementary Figure 3.**
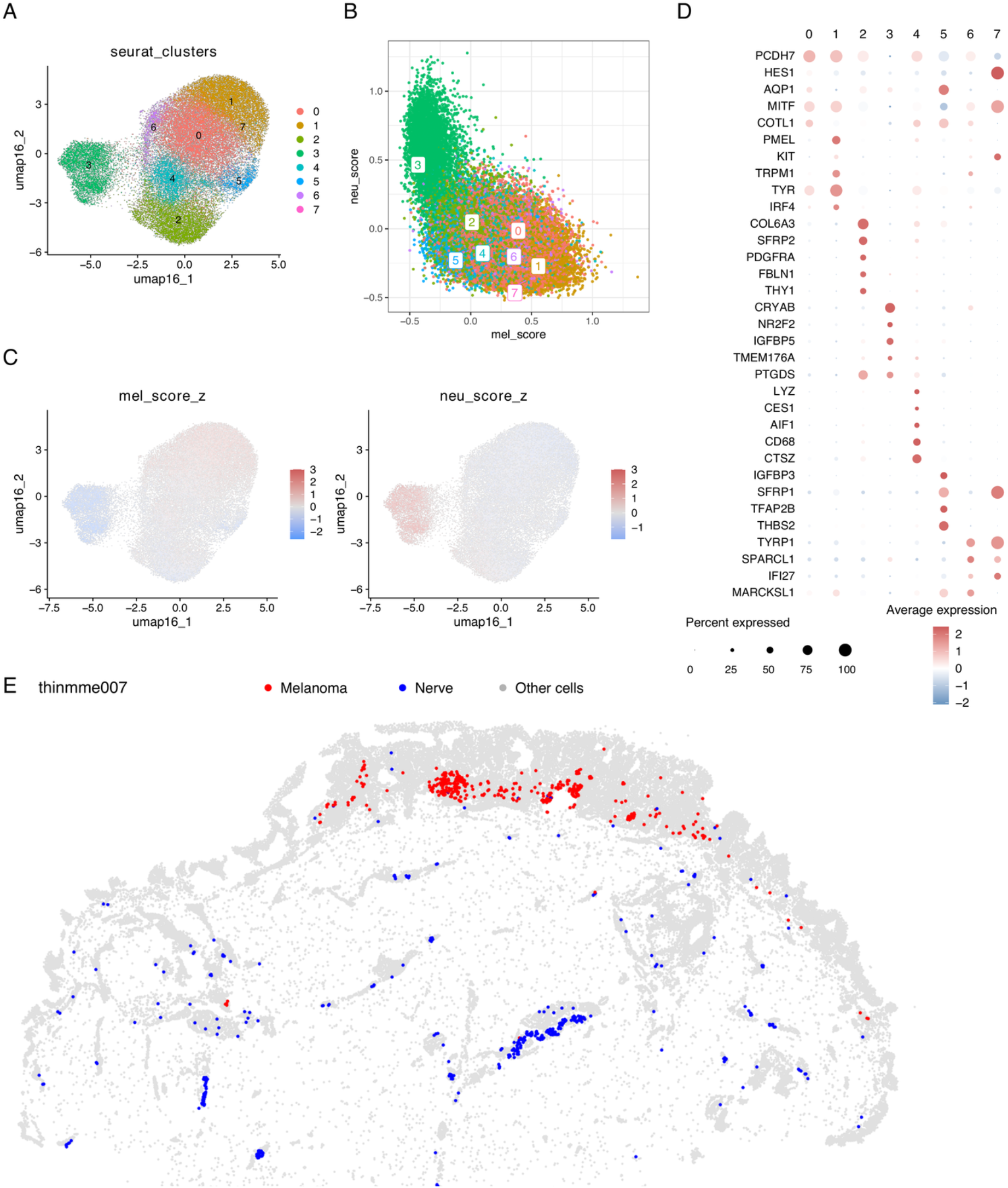
Resolution of mixed melanoma/nerve cluster 16. **A)** UMAP embedding of re-analyzed cluster-16 cells, colored by de novo Seurat subcluster identity. **B)** Scatter plot of melanoma and neural module scores for each cell. Subcluster labels indicate median score positions. Melanoma scores were computed from PMEL, TYR, S100A13, MLANA, MITF, TYRP1, PCDH7, QPCT, EDNRB, and MYC; neural scores from CRYAB, IGFBP5, NR2F2, TMEM176A, ITGA6, PLP1, DST, PTGDS, CXCL12, and MYL9. **C)** UMAPs colored by z-scored melanoma and neural module scores. **D)** Dot plot showing marker expression across cluster-16 subclusters. Dot size indicates the percentage of cells expressing each marker and color indicates scaled average expression. **E)** Representative spatial reconstruction showing classified cluster-16 cells mapped back to tissue coordinates. Cells assigned as melanoma are shown in red, nerve-associated cells in blue, and all other cells in grey.

**Supplementary Figure 4.**
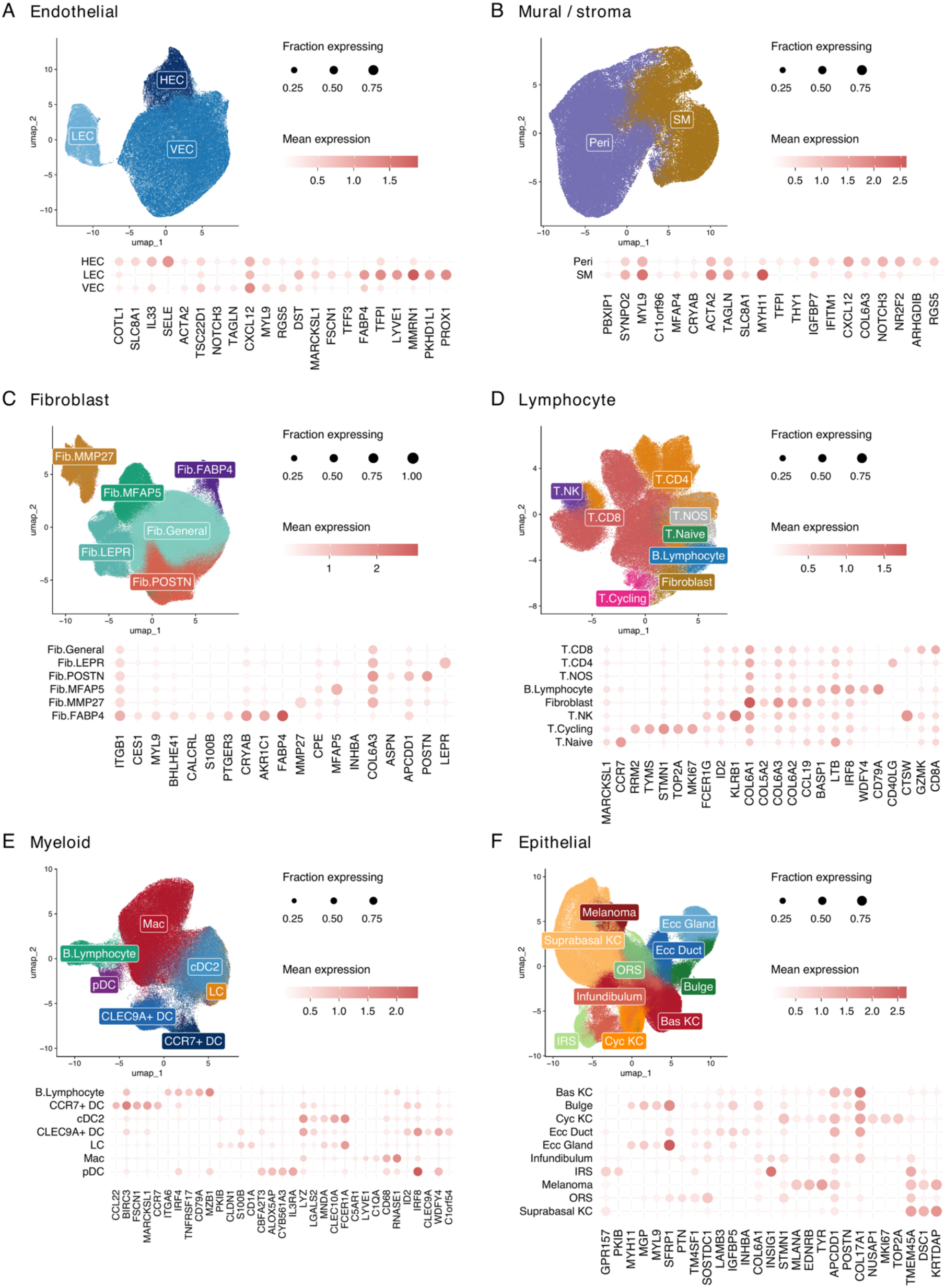
Subtype annotation of major non-melanoma cellular compartments in the Xenium dataset. Annotated UMAPs and corresponding subtype-marker dot plots for **(A)** endothelial, **(B)** mural/stromal, **(C)** fibroblast, **(D)** lymphocyte, **(E)** myeloid and **(F)** epithelial compartments. Dot size indicates the fraction of cells expressing each gene, and color indicates mean expression, using compartment-specific color scales.

**Supplementary Figure 5.**
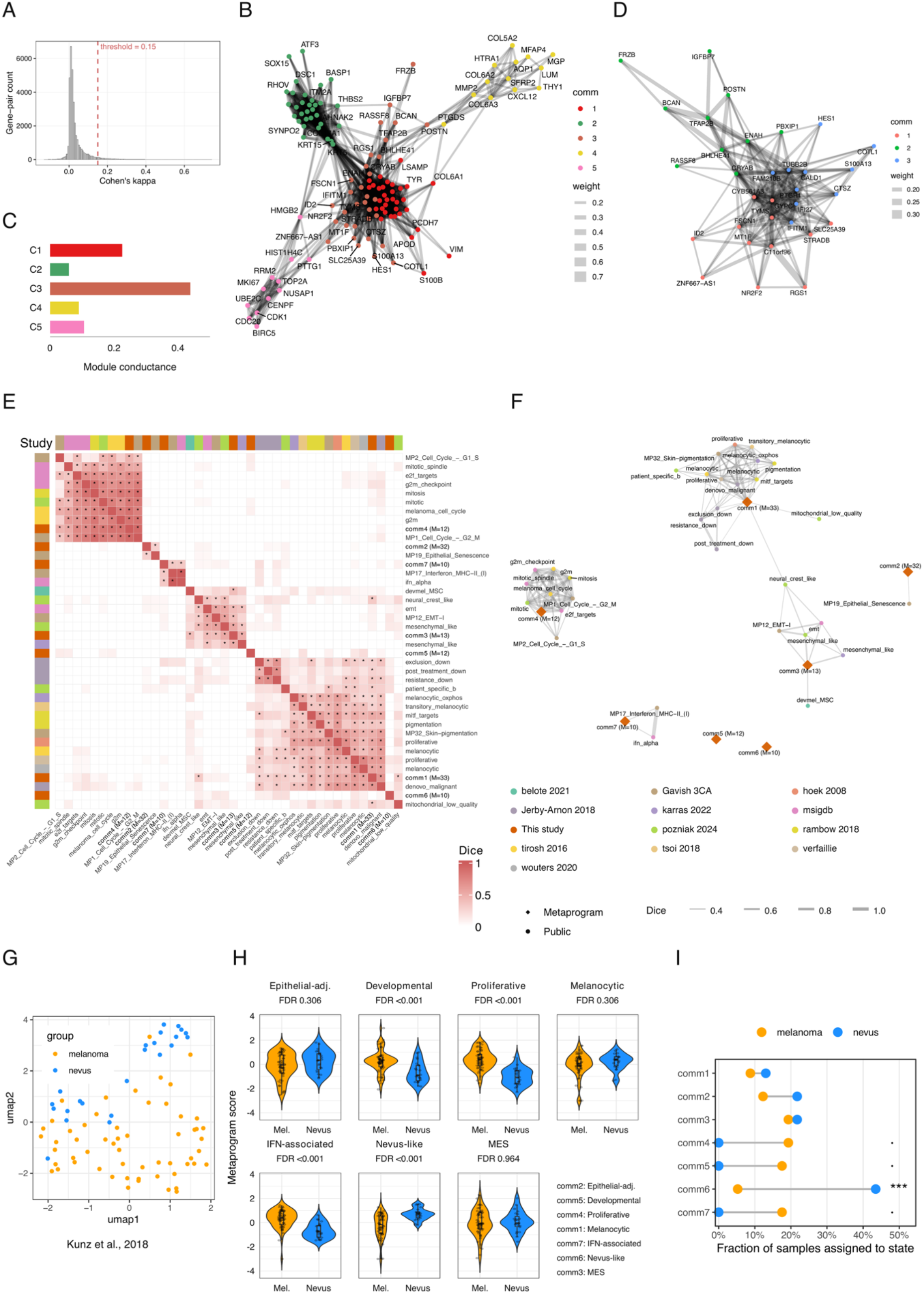
Identification and annotation of melanoma metaprograms. **A)** Distribution of pairwise gene co-detection across melanoma cells, quantified using Cohen’s κ on binarized detection. The dashed line marks the retention threshold (κ = 0.15). **B)** Gene co-detection network retained at κ > 0.15, restricted to its largest connected component and genes of degree > 1. Colors indicate the five modules identified by Louvain clustering. **C)** Conductance of the initial modules. Module 3 had the highest conductance (0.44) and was selected for further subdivision. **D)** Re-clustering of module 3 into three subprograms, yielding seven final metaprograms across the 122 network genes. **E)** Pairwise Dice similarity including the seven final metaprograms and retained public signatures, calculated using genes represented in the Xenium panel. Asterisks denote hypergeometric enrichment at FDR < 0.10, with Benjamini–Hochberg correction. **F)** Network representation of these associations with public signatures. Edges denote associations passing the threshold, with width proportional to Dice similarity. M indicates the number of genes in each metaprogram. **G)** UMAP of the independent bulk transcriptomic cohort of melanocytic nevi and primary melanomas (Kunz et al., 2018), calculated from the seven metaprogram scores. **H)** Continuous metaprogram scores in melanomas and nevi. FDR values are Benjamini–Hochberg corrected across the seven metaprograms. **I)** Fraction of expression profiles assigned to each dominant metaprogram, defined by the highest standardized score. Group differences were assessed using two-sided Fisher’s exact tests comparing each metaprogram against all others, with Benjamini–Hochberg correction across the seven tests (· FDR < 0.10; *** FDR < 0.001).

**Supplementary Figure 6.**
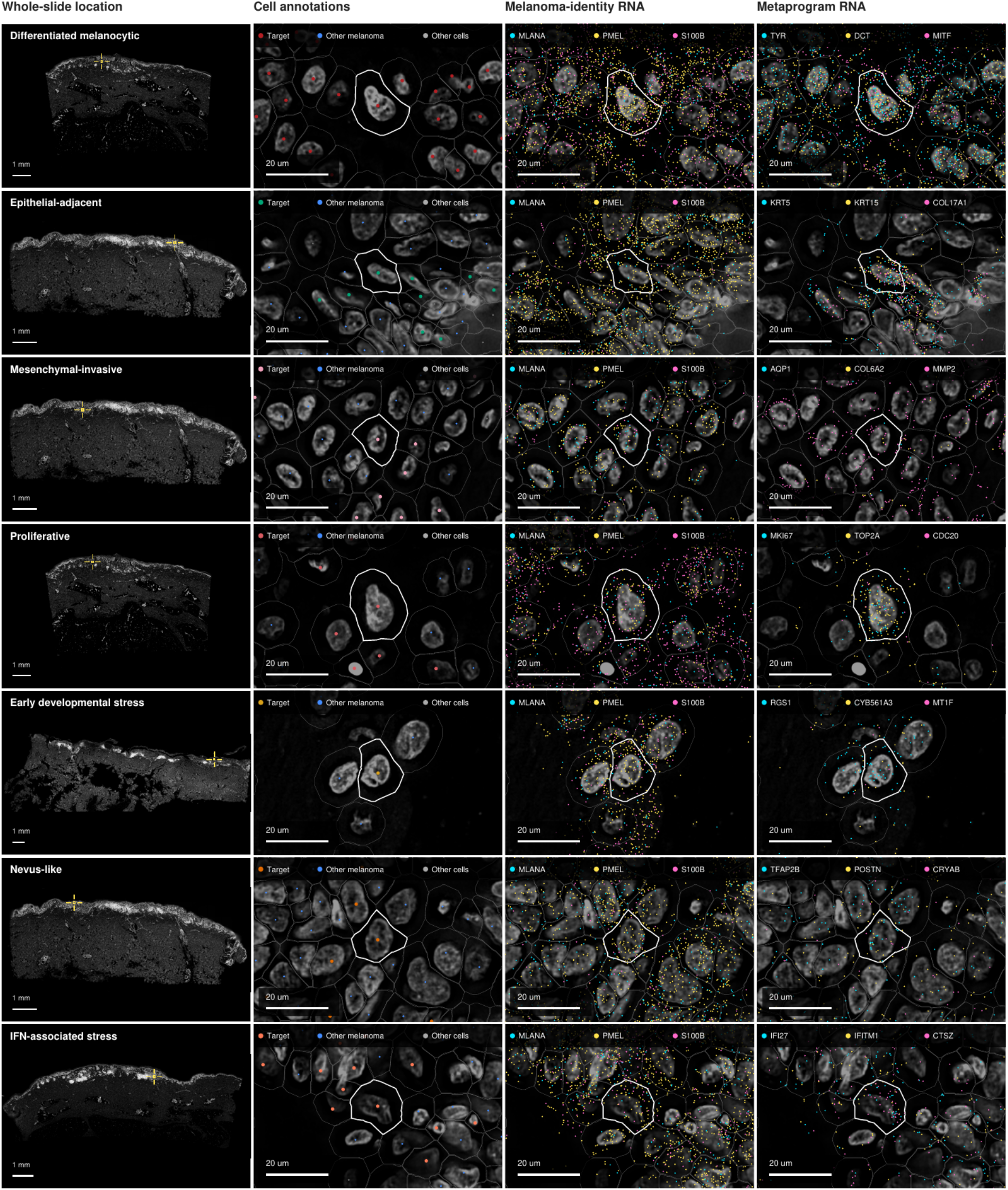
Spatial transcript examples of the seven melanoma metaprograms. Each row shows a selected example of the indicated metaprogram. Columns show the whole-section location of the selected region, cell annotations, melanoma-associated transcripts (MLANA, PMEL and S100B), and representative metaprogram transcripts. Enlarged views show the same region overlaid on DAPI. Target cells are colored by metaprogram, other melanoma cells in blue and non-melanoma cells in grey. Segmentation boundaries are shown in grey, with the selected cell outlined in white.

**Supplementary Figure 7.**
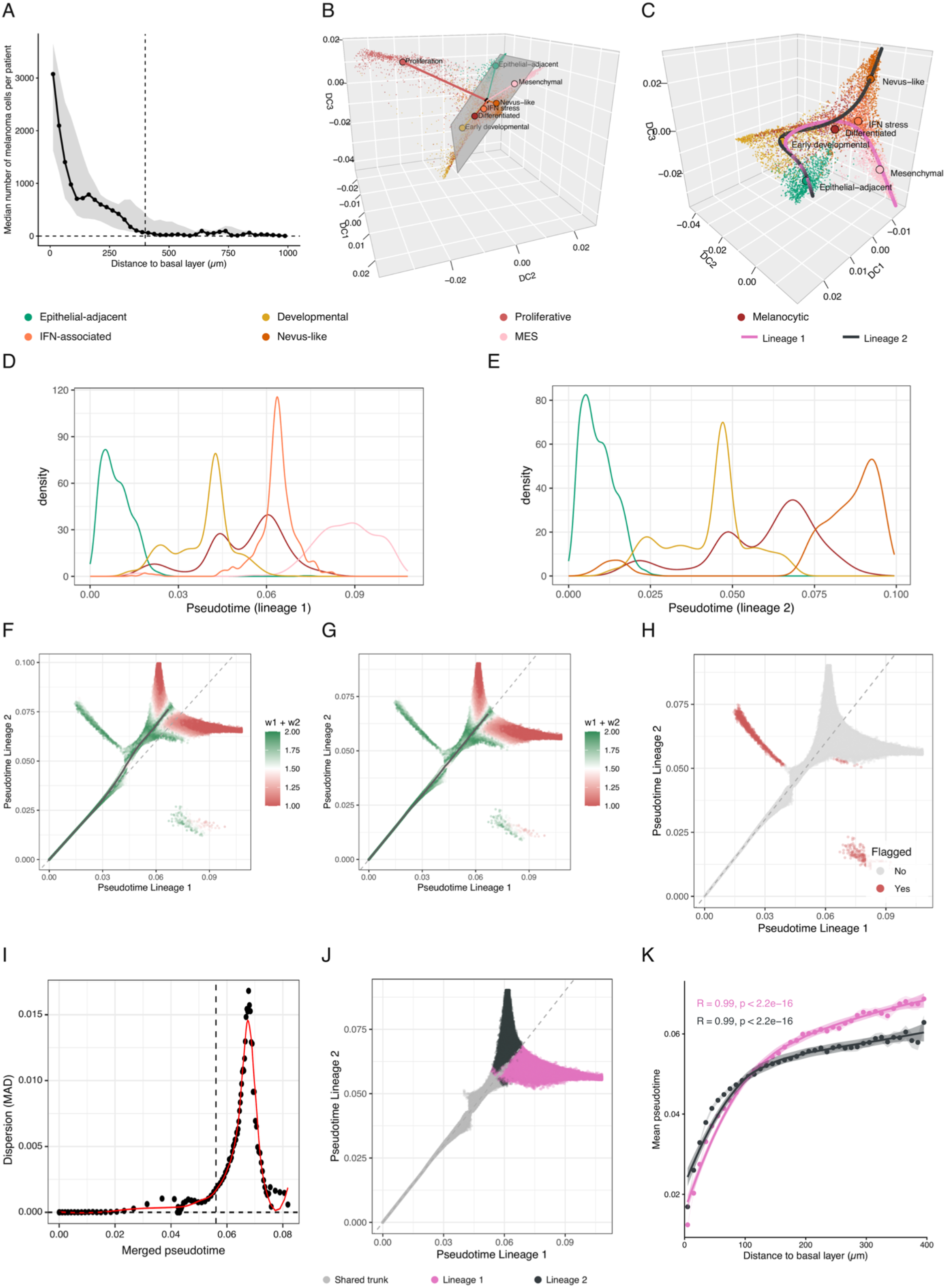
Trajectory inference of melanoma cell states. **A)** Melanoma-cell abundance in 25-µm depth bins relative to the epidermal basal layer. Points show median counts across patients represented in each bin, shading indicates the interquartile range. The dashed vertical line marks the 400-µm range used for subsequent analyses. **B)** Three-dimensional diffusion-map representation of melanoma cells colored by metaprogram. A plane was fitted to the centroids of the six non-proliferative metaprograms; the proliferation/cell-cycle cells extend almost perpendicularly to this plane. **C)** Diffusion-map embedding constructed after excluding the proliferation program, with the two inferred Slingshot trajectories overlaid. **D–E)** Kernel-density distributions of metaprogram-labeled reference cells along Lineage 1 (D) and Lineage 2 (E) pseudotime, illustrating their ordering along each trajectory. **F–G)** Relationship between the two lineage pseudotimes after projection of the complete melanoma dataset, colored by the sum of lineage-assignment weights, w1+w2, before (F) and after (G) alignment of pseudotime values along the shared trunk. **H)** Identification of cells lying far from their assigned trajectory centerline. **I)** Dispersion of the difference between lineage pseudotimes along the merged progression axis. The red curve shows the smoothed trend, and the vertical dashed line indicates the selected fork for branch divergence. **J)** Final classification of melanoma cells into shared trunk, Lineage 1 or Lineage 2 after pseudotime alignment and reassignment of ffagged cells. **K)** Relationship between tissue depth and lineage pseudotime. Points represent mean pseudotime across melanoma cells pooled within 10-µm depth bins, curves show smoothed trends, with shading indicating 95% confidence intervals for bin means and fitted trends. Spearman correlations are displayed.

**Supplementary Figure 8.**
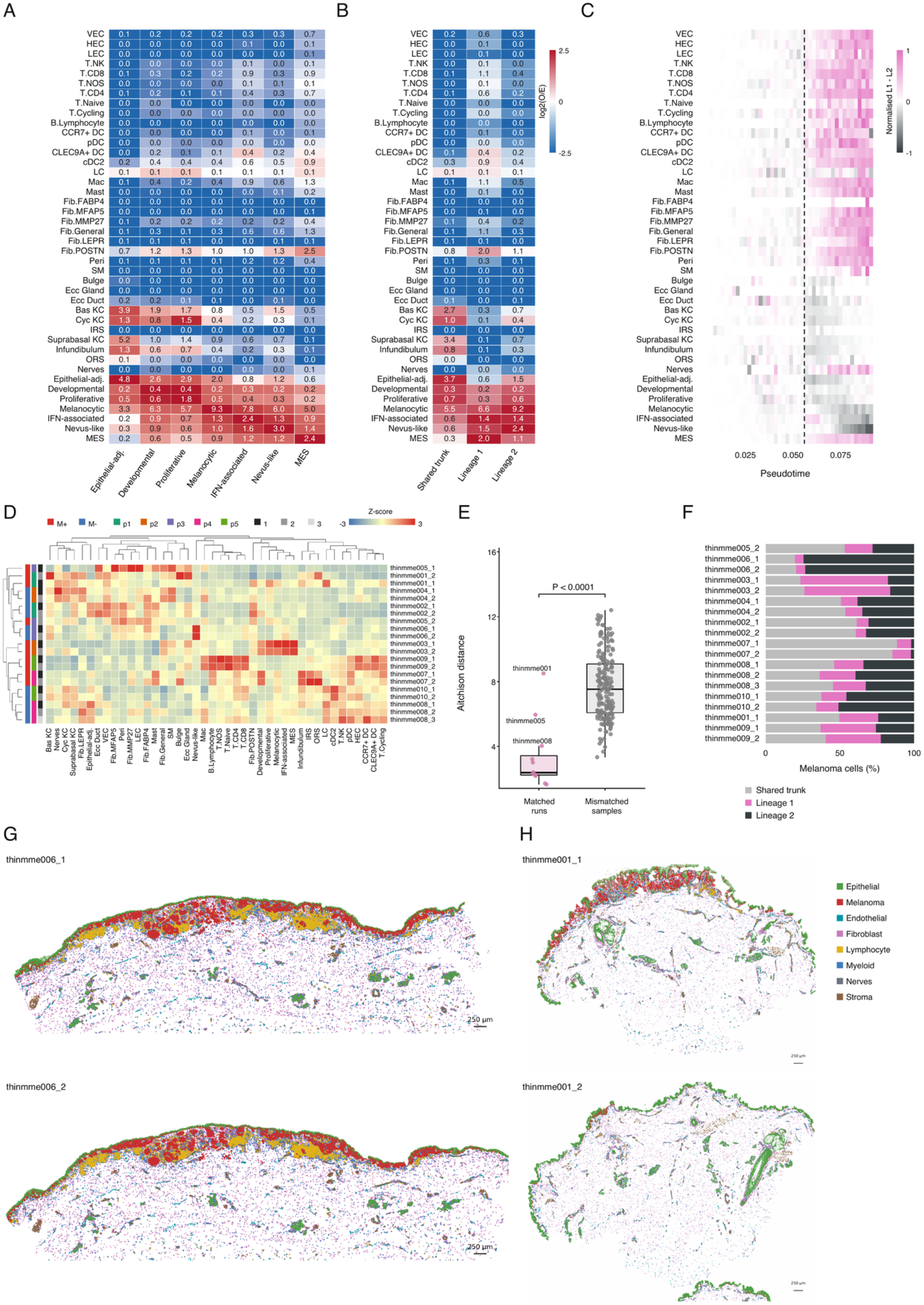
Neighborhood organization and repeat-section concordance. **A–B)** Composition of the 25 nearest neighbors of melanoma cells, grouped by metaprogram (A) or trajectory class (B). Numbers indicate the median across acquisitions of the mean neighbor count per reference cell. **C)** Neighborhood differences between Lineage 1 and Lineage 2 along pseudotime. Differences were normalized by the maximum neighbor proportion across both lineages and displayed pseudotime bins, separately for each neighbor type. The dashed line marks the inferred fork. D) Hierarchically clustered heatmap of cell-state composition across all 21 Xenium acquisitions. Z-scores were calculated per cell type. **E)** Aitchison distances between cell-composition profiles from repeated acquisitions of the same tumor and acquisitions from different tumors. The permutation test assessed whether repeated acquisitions were compositionally closer than expected under random sample matching. **F)** Proportions of melanoma cells assigned to the shared trunk, Lineage 1 or Lineage 2 in each of the 19 retained acquisitions. **G–H)** Representative paired tissue reconstructions from thinmme006 (G) and thinmme001 (H), colored by broad cellular lineage.

**Supplementary Figure 9.**
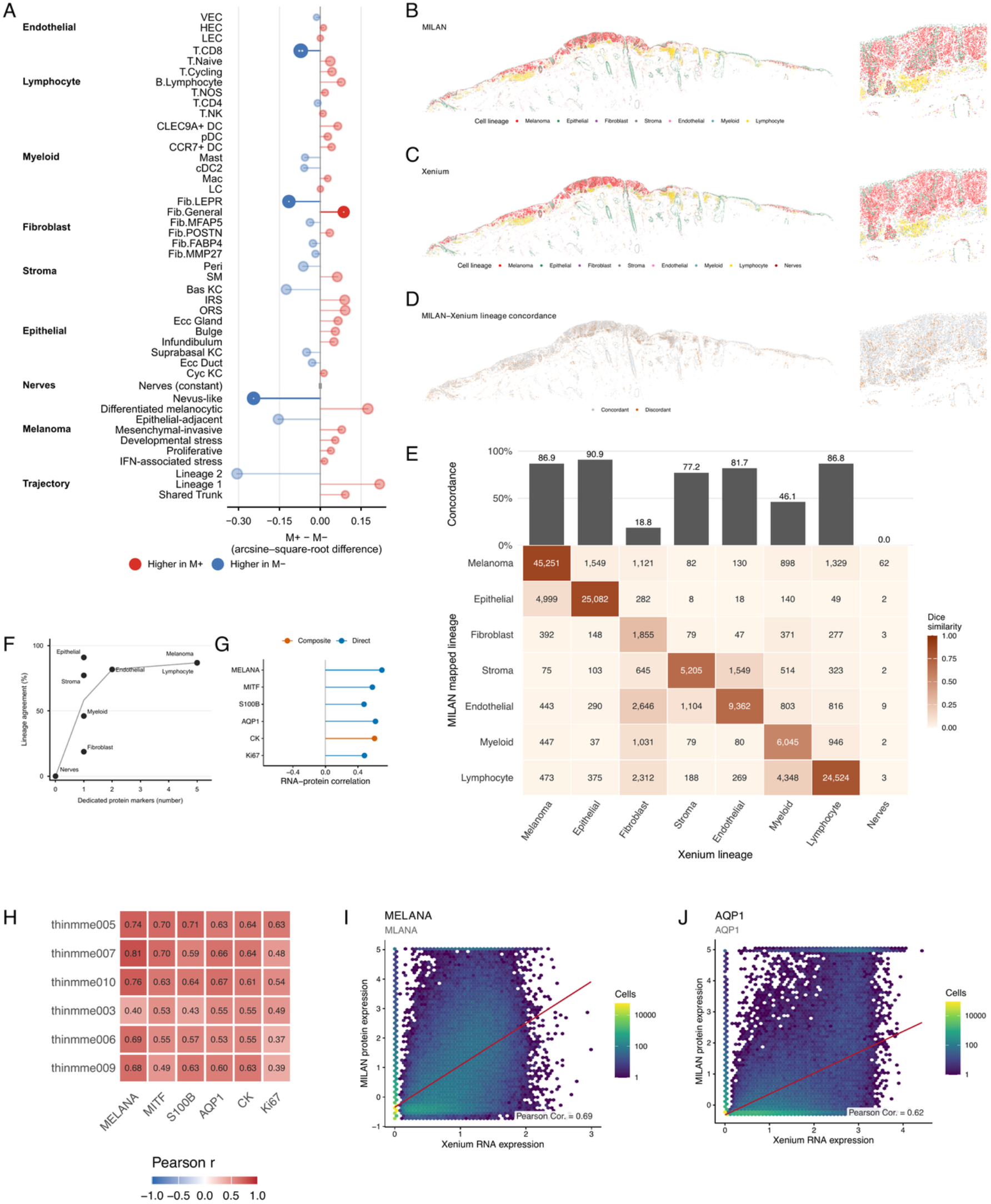
Cellular composition and cross-modal RNA–protein concordance. **A)** Paired propeller analysis of patient-level cell proportions within each broad cellular compartment. Effects represent M+ minus M− differences on the arcsine–square-root scale. Point size increases with −log₁₀(FDR); asterisks indicate Benjamini– Hochberg-adjusted significance within each compartment (* FDR < 0.05; ** FDR < 0.01). **B–D)** Registered tissue maps and corresponding close-ups of matched cells in thinmme007, colored by independently derived MILAN annotations (B), Xenium annotations (C), or agreement between their mapped lineage labels (D). **E)** Lineage concordance excluding MILAN Blank and Noise assignments. Bars show the percentage of matched cells within each Xenium lineage. Heatmap numbers indicate matched-cell counts. **F)** Relationship between the number of dedicated MILAN markers and lineage concordance. **G–H)** Pearson RNA–protein correlations across pooled matched cells (G) and separately for each sample (H). Direct comparisons use the corresponding transcript; the composite CK comparison uses mean expression of KRT5, KRT15 and KRT18. **I–J)** Joint distributions of normalized RNA and protein measurements for MLANA/MELANA (I) and AQP1 (J). Red lines indicate linear fits.

**Supplementary Figure 10.**
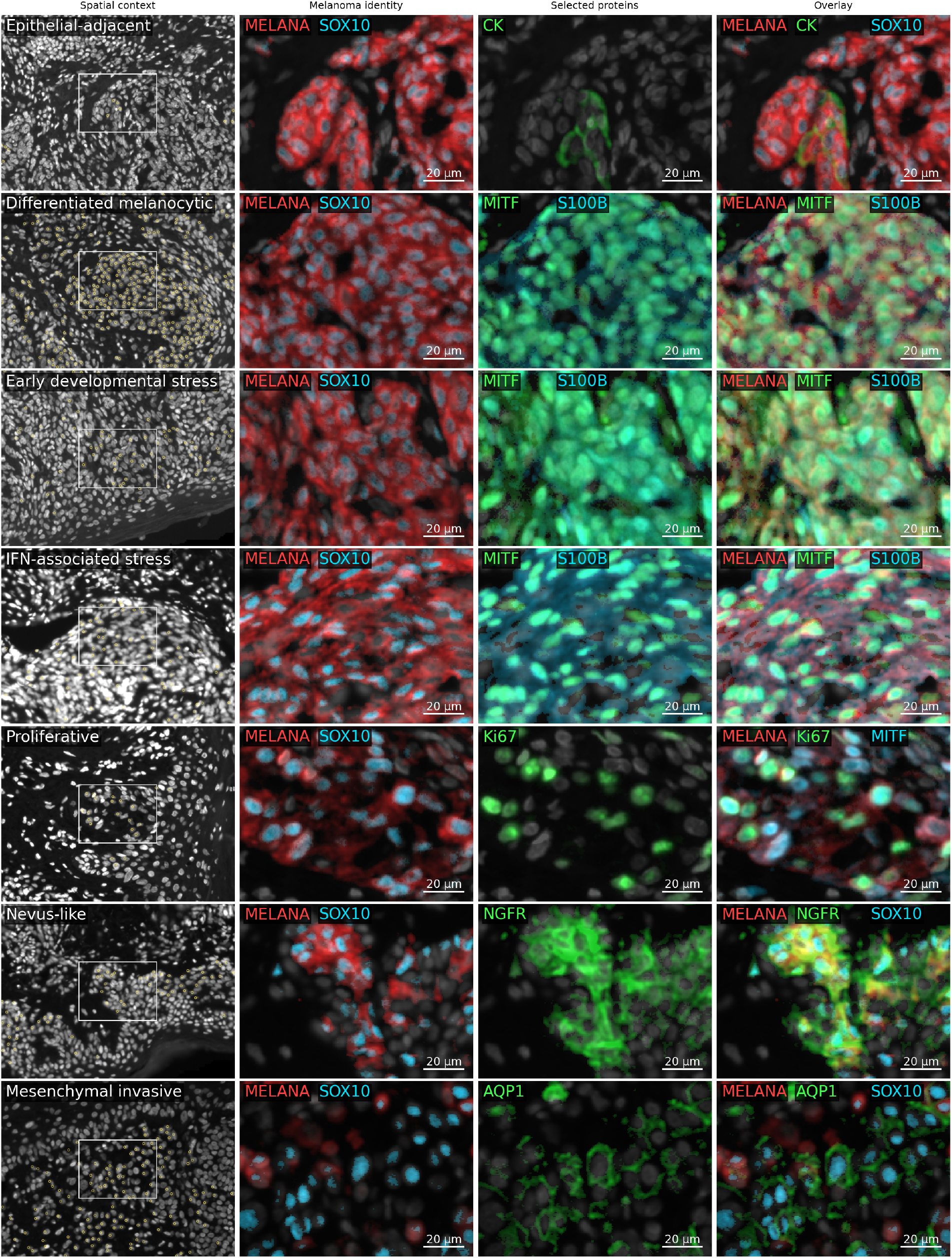
Protein-expression examples of Xenium-defined melanoma metaprograms. Multiplex immunoffuorescence images of representative regions containing cells assigned to the seven Xenium-defined melanoma metaprograms. The first column shows a DAPI staining, with yellow points marking matched cells assigned to the indicated metaprogram and white boxes identifying the enlarged regions. Subsequent columns show MELANA/SOX10 as melanoma lineage protein markers, and selected proteins markers for each metaprogram, and combined overlays. Developmental and IFN-associated rows illustrate melanocytic protein expression rather than state-specific protein markers.

**Supplementary Figure 11.**
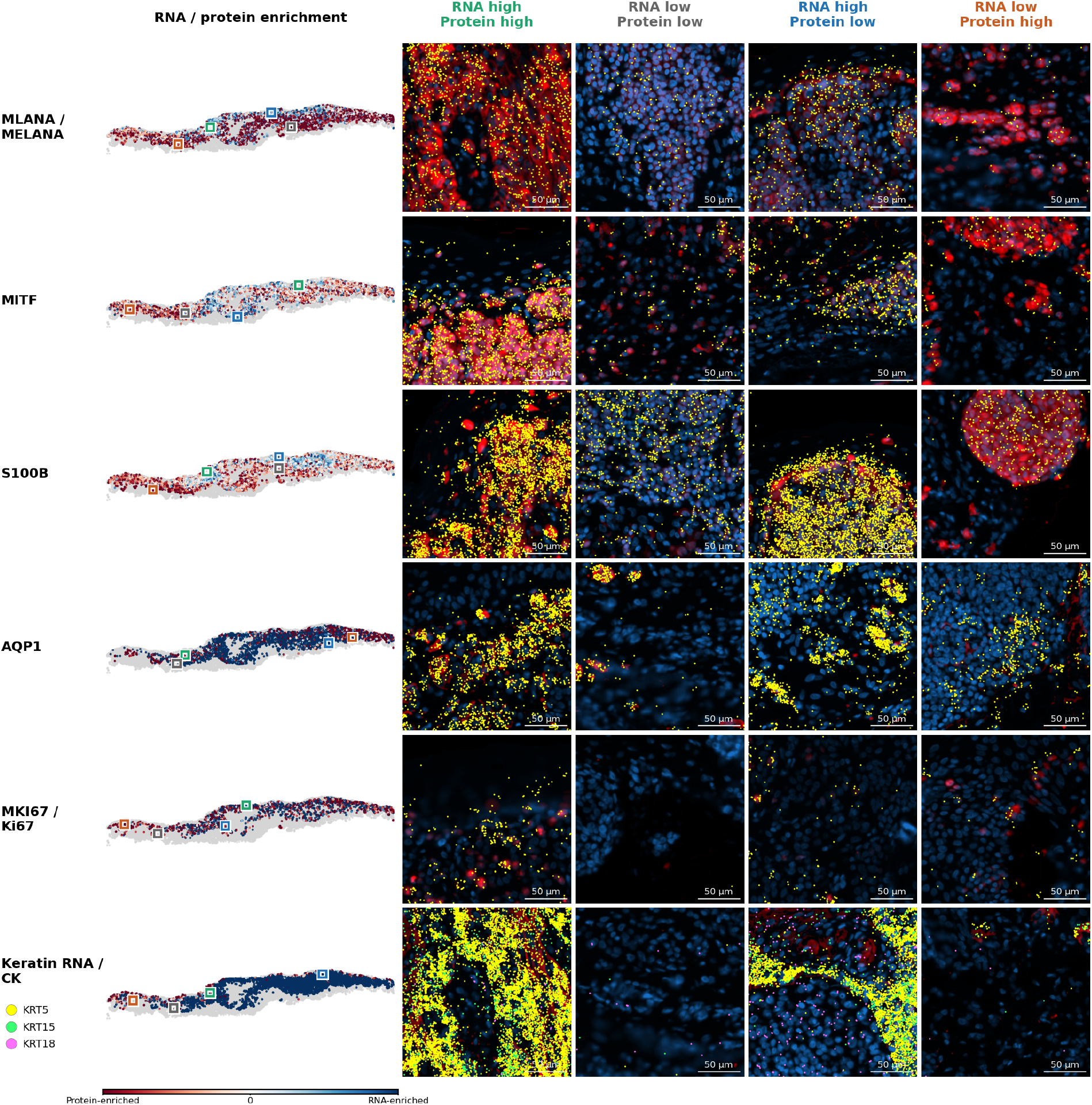
Spatial comparison of RNA and protein abundance in melanoma. Same-section Xenium and MILAN measurements in thinmme003 are shown for MLANA/MELANA, MITF, S100B, AQP1, MKI67/Ki67 and keratin RNA/CK. The first column shows relative RNA-to-protein enrichment in matched melanoma cells, with blue indicating RNA enrichment and red indicating protein enrichment. Overlays show DAPI in blue, protein staining in red and individual Xenium transcripts in yellow. Four snapshots were selected: protein-high rna-high, protein-high rna-low, protein-low rna-high, and protein-low rna-low.

**Supplementary Figure 12.**
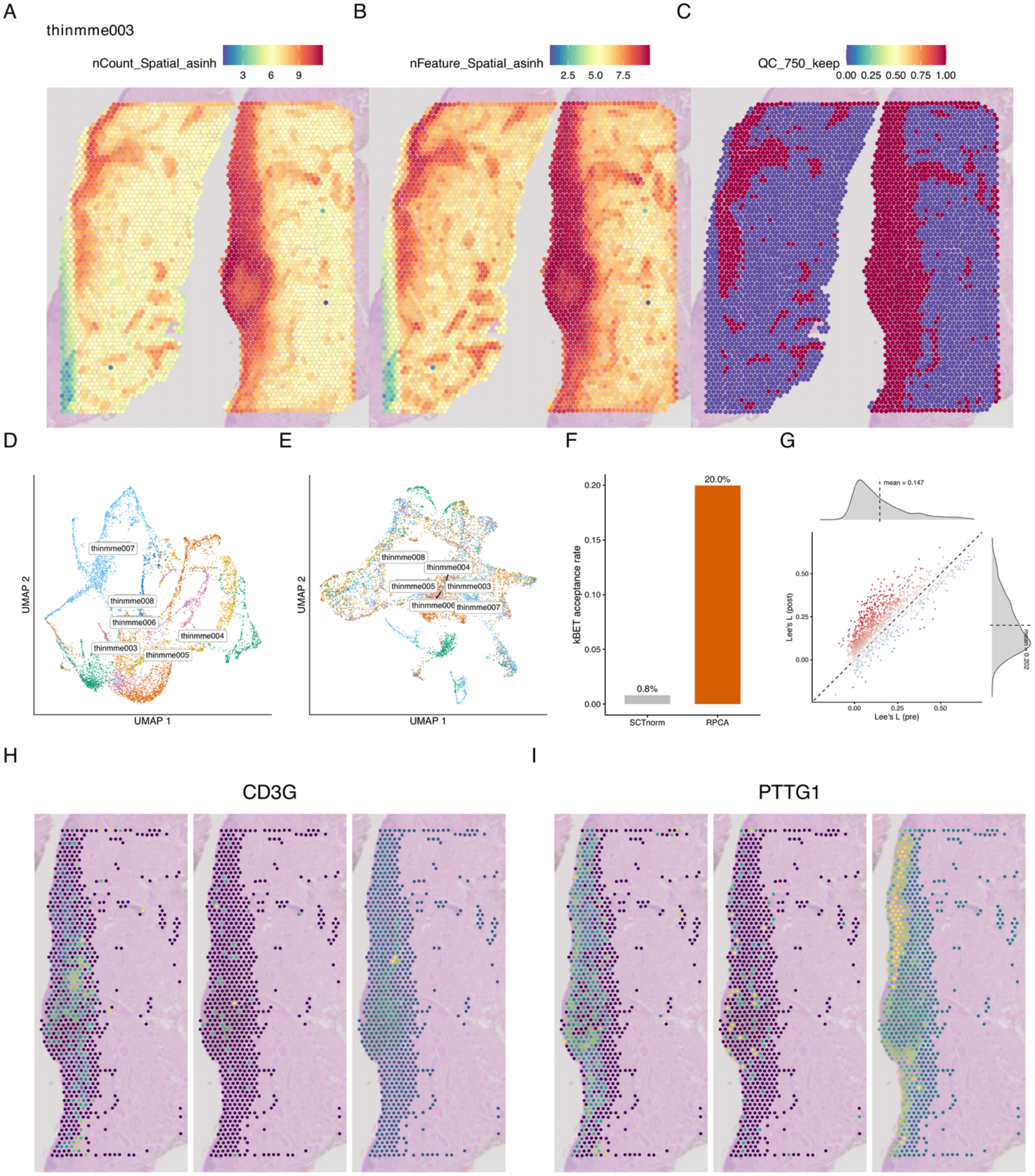
Quality control, normalization, integration and imputation of Visium sections. **A–C)** Representative tissue section (thinmme003) showing total transcript counts (asinh scale), the number of detected genes (asinh scale), and whether spots passed the QC threshold of ≥750 detected genes, overlaid on the H&E image. D–E) Two-dimensional UMAP representations of section-wise SCTransform-normalized data before (D) and after (E) reciprocal principal-component analysis (RPCA) integration, colored by tissue section. F) Cross-section mixing quantified by the k-nearest-neighbor batch-effect test (kBET) acceptance rate. Neighborhoods comprised 30 spots in the first five principal components; acceptance rates represent the mean proportion of neighborhoods without a significant deviation from the expected section composition (p ≥ 0.05), across 100 repeats. G) Gene-wise spatial correspondence between registered Xenium pseudospots and Visium expression, measured using Lee’s L before (x-axis) and after (y-axis) MAGIC imputation. The dashed diagonal indicates equal correspondence before and after imputation. H–I) Spatial expression of CD3G (H) and PTTG1 (I), showing registered Xenium pseudospots, pre-MAGIC SCTransform-normalized Visium expression, and post-MAGIC expression from left to right.

**Supplementary Figure 13.**
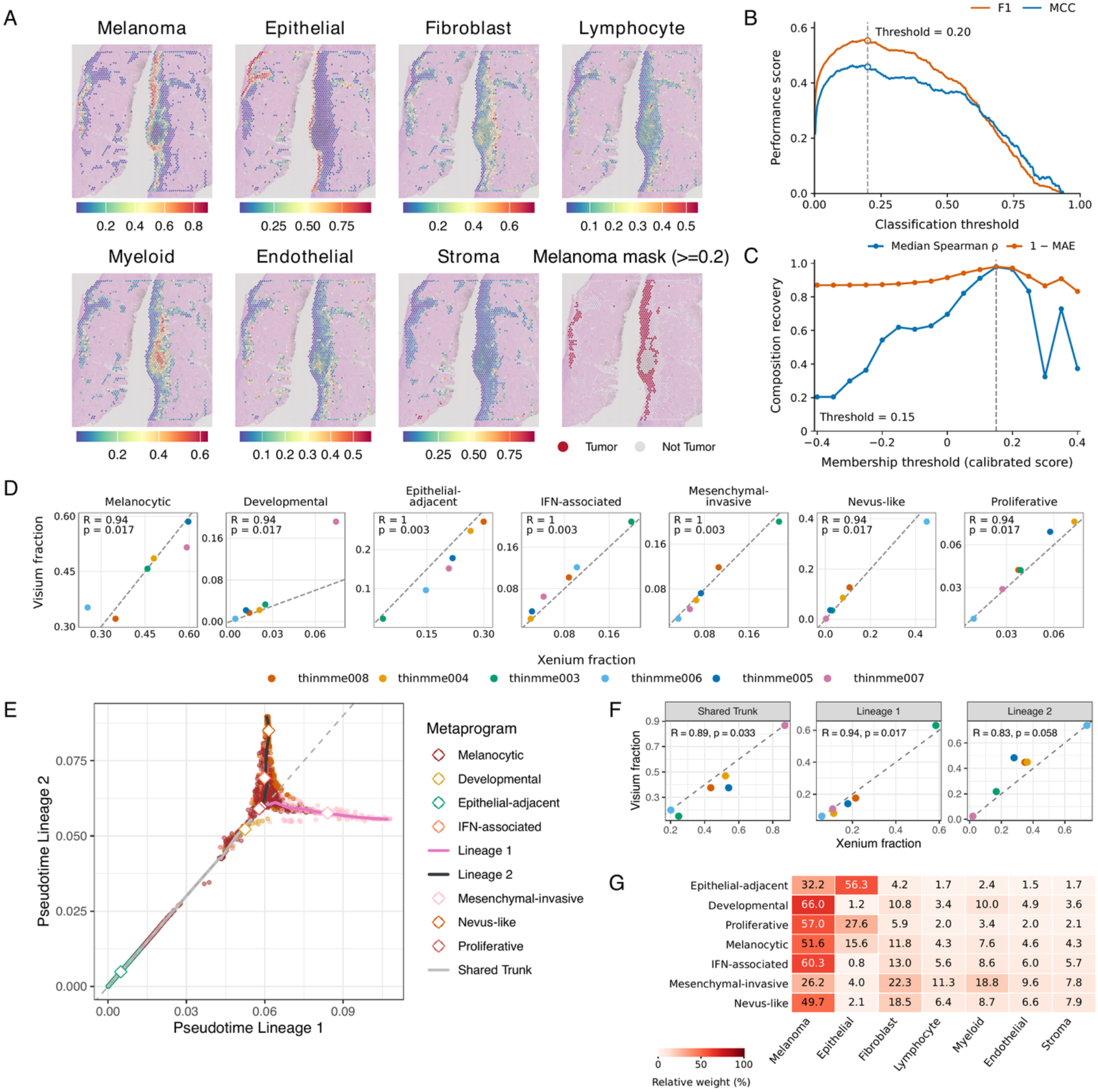
Xenium-guided deconvolution and projection of melanoma metaprograms and trajectories onto Visium. **A)** Relative cell2location weights for the main cell types and the selected melanoma-associated spot mask, overlaid on H&E for thinmme003. **B)** F1 and MCC scores across melanoma-weight thresholds, evaluated against registered Xenium pseudospots. The dashed line indicates the selected threshold of 0.20. **C)** Median within-patient Spearman correlation and composition agreement (1-MAE) between Visium and Xenium metaprogram compositions. The dashed line indicates the selected threshold of 0.15. **D)** Patient-level correspondence between Xenium melanoma-cell fractions and Visium metaprogram composition. **E)** Visium spots (n = 1,590) in projected Lineage 1–Lineage 2 pseudotime space, colored by dominant metaprogram. **F)** Correspondence between patient-level Xenium melanoma-cell and Visium-spot fractions assigned to the shared trunk, Lineage 1 or Lineage 2. **G)** Mean relative cell2location weights (%) within each metaprogram-defined spot group, showing their estimated cellular composition.

